# Evolution of circadian behavioral plasticity through *cis*-regulatory divergence of a neuropeptide gene

**DOI:** 10.1101/2023.07.05.547553

**Authors:** Michael P. Shahandeh, Liliane Abuin, Lou Lescuyer De Decker, Julien Cergneux, Rafael Koch, Emi Nagoshi, Richard Benton

## Abstract

Widely-distributed species experience substantial environmental variation, which they often accommodate through behavioral plasticity. Although this ability is integral to fitness, we have little understanding of the mechanistic basis by which plasticity evolves. One factor that varies seasonally and by latitude is photoperiod (day length). Many organisms, including the cosmopolitan *Drosophila melanogaster* display circadian plasticity, adjusting to fluctuating photoperiod by varying the timing of their activity to coincide with changing dawn/dusk intervals^1^. Here, we compare *D. melanogaster* with the closely-related ecological specialist *Drosophila sechellia*, an equatorial island endemic that experiences minimal photoperiod variation, to investigate the molecular-genetic basis of circadian plasticity evolution^2,3^. We discover that *D. sechellia* displays exceptionally little circadian plasticity compared to *D. melanogaster* and other non-equatorial drosophilids. Through a screen of circadian mutants in *D. melanogaster*/*D. sechellia* hybrids, we identify a role of the neuropeptide Pigment-dispersing factor (Pdf) in this loss. While the coding sequence of *Pdf* is conserved, we show that *Pdf* has undergone *cis*-regulatory divergence in these drosophilids. We document species-specific temporal dynamic properties of *Pdf* RNA and protein expression, as well as Pdf neuron morphological plasticity, and demonstrate that modulating Pdf expression in *D. melanogaster* can influence the degree of behavioral plasticity. Furthermore, we find that the *Pdf* regulatory region exhibits signals of selection across populations of *D. melanogaster* from different latitudes. Finally, we provide evidence that plasticity confers a selective advantage for *D. melanogaster* at higher latitudes, while *D. sechellia* likely suffers fitness costs through reduced copulation success outside its range. Our work defines *Pdf* as a locus of evolution for circadian plasticity, which might have contributed to both *D. melanogaster*’s global distribution and *D. sechellia*’s habitat specialization. Moreover, together with spatial changes in Pdf expression reported in high-latitude drosophilid species^4,5^, our findings highlight this neuropeptide gene as a hotspot for circadian plasticity evolution.

## Introduction

Nervous systems coordinate animals’ behavioral responses to the external world to maximize survival and fitness. This task becomes more challenging when environments are not constant, a problem of substantial significance for broadly-distributed species. One way to face changing conditions is with behavioral plasticity, that is, the ability to adjust behavioral phenotypes to match fluctuations in the environment. There are many examples of plastic behaviors in nature: songbirds shift the frequency of their vocalizations in response to anthropogenic noise^6^, ants alter their locomotor and foraging behaviors as a function of temperature^7^, and lizards change their basking behavior based on altitude^8^. However, we have little understanding of whether and how behavioral plasticity is determined and evolves at the genetic and cellular level.

An important example of plastic behavior in animals is circadian activity, whereby species adjust their daily activity patterns in response to seasonal variation in day length^9^. This ability is critical because circadian activity in most animals coordinates specific behaviors with optimal activity periods throughout the day to, for example, avoid environmental stressors, maximize food availability, and align with conspecifics for synchronized social and sexual behaviors^10,11^. As such, deviations from regular circadian patterns can negatively affect fitness and species persistence^12,13^. Drosophilids are a powerful system to study circadian behavioral plasticity. These flies display large bouts of activity surrounding dawn and dusk (termed morning and evening activity peaks), separated by a period of relative inactivity during the middle of the day^14^. The best-studied species, the cosmopolitan *Drosophila melanogaster*, plasticly adjusts its circadian rhythm depending upon seasonal variation in photoperiod^1^. Notably, the degree of photoperiod plasticity of different strains of this species correlates with the latitude of collection site^15^. Moreover, several distantly-related, high-latitude species have evolved divergent patterns of activity and extreme plasticity, allowing their daily activity to match long summer days^16^.

A potentially interesting comparison species to *D. melanogaster* is *Drosophila sechellia*, a much closer relative that diverged 3-5 million years ago (Fig. 1a)^2,3^. *D. sechellia* is endemic to the equatorial islands of the Seychelles archipelago, where it experiences almost no seasonal photoperiod variation (Fig. 1a-b). Here, we discover striking differences in the circadian activity and plasticity of *D. sechellia* and *D. melanogaster*, notably an almost complete inability of *D. sechellia* to adapt to increased photoperiod. Taking advantage of the possibility to interbreed these species, we conducted a genetic screen of known circadian genes, offering unprecedented insights into the genetic and cellular underpinnings of circadian plasticity evolution, and a rare connection between genetic divergence and evolved differences in an ecologically-relevant behavior.

**Fig. 1.**
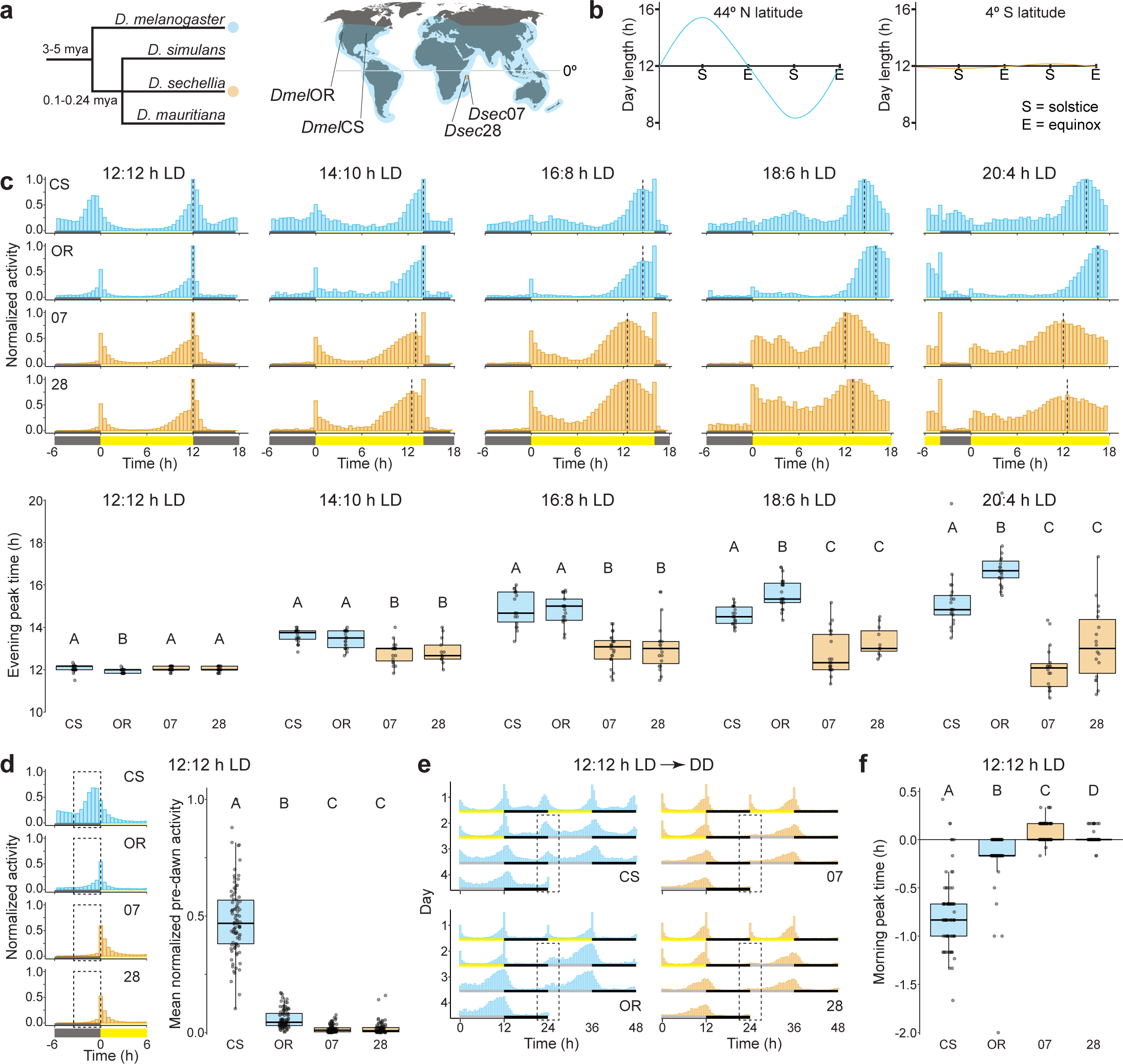
*D. sechellia* displays reduced circadian plasticity and lower morning activity than *D. melanogaster*. **a**, Left: phylogeny of the *Drosophila melanogaster* sub-group. Right: modern ranges of the focal species of this study, *D. melanogaster* (*Dmel*) and *D. sechellia* (*Dsec*), are indicated by the shaded regions (blue and orange, respectively) on the map, with the approximate collection sites of the wild-type strains used. **b**, Approximate seasonal photoperiod variation at the collection sites of the *D. melanogaster* (left) and *D. sechellia* strains (right). **c**, Top: mean normalized activity of two *D. melanogaster* (CS and OR, blue) and two *D. sechellia* (07 and 28, orange) strains under the indicated photoperiods. Plots depict normalized average activity of the last 4 days of a 7-day photoperiod, for extended photoperiods, following 7 days of 12:12 h LD. Vertical dashed lines indicate the average timing of the evening peak for each strain. Here and elsewhere, yellow and grey bars indicate timing of lights-on and lights-off, respectively. Overall, *D. sechellia* strains were slightly less active than *D. melanogaster* strains. Bottom: box plots depict evening peak time quantifications for individual flies under each photoperiod. Here and elsewhere, box plots show the median (bold line), interquartile range (box), and whiskers represent the final quartiles. All data points are shown overlaid on box plots. Outliers are points that fall beyond the box plot whiskers. Letters indicate significant differences: p < 0.05 (pairwise Wilcoxon test with Bonferroni correction). Sample sizes (numbers of individual flies) are as follows: 12:12 h LD: CS (18), OR (21), 07 (24), 28 (19); 14:10 h LD: CS (22), OR (22), 07 (19), 28 (13); 16:8 h LD: CS (18), OR (21), 07 (24), 28 (19); 18:10 h LD: CS (22), OR (23), 07 (21), 28 (11); 20:4 h LD: CS (21), OR (22), 07 (19), 28 (18). **d**, Mean normalized activity of *D. melanogaster* and *D. sechellia* strains under a 12:12 h LD cycle during the morning activity peak (same data from **c.** restricted to -6 to 6 h). Left: plots depict average activity of the last 4 days of a 7-day recording period. Dashed boxes highlight the pre-dawn period, 3 h before lights-on. Right: mean normalized activity of individual flies within this pre-dawn period. Letters indicate significant differences: p < 0.001 (pairwise Wilcoxon test with Bonferroni correction). Sample sizes as follows: CS (89), OR (93), 07 (95), 28 (91). **e**, Double plotted actograms depicting the transition from the last 2 days of 12:12 h LD to the first 2 days of constant darkness (DD) for each strain. Dashed boxes highlight morning activity peak period during DD, 3 h before and after subjective lights-on. Sample sizes as follows: CS (29), OR (32), 07 (29), 28 (22). Grey bars indicate timing of subjective lights-on during DD. **f**, Morning peak time, in hours from lights-on, for individual flies from **d**. Letters indicate significant differences: p < 0.001 (pairwise Wilcoxon test with Bonferroni correction).

## Results

### *D. sechellia* displays reduced photoperiod plasticity compared to *D. melanogaster*

To test for species-specific differences in circadian plasticity, we first measured circadian behavior for *D. melanogaster* and *D. sechellia* under a standard 12 h light-dark cycle (12:12 h LD) as well as four extended photoperiod regimes ranging from mild (14:10 h LD) to extreme (20:4 h LD) (Fig. 1c). We used males of two strains each of *D. melanogaster* and *D. sechellia* (Supplementary Table 1), to distinguish interspecific from intraspecific phenotypic differences. The two *D. melanogaster* strains (*Dmel*CS and *Dmel*OR) were collected at ∼41° N and ∼44° N, respectively, while the *D. sechellia* strains (*Dsec*07 and *Dsec*28) are from the Seychelles archipelago, at ∼4° S of the equator (Fig. 1a). The strains of each species therefore initially evolved in environments where they experienced large differences in annual photoperiod variation (Fig. 1b). Under each photoperiod, all strains displayed activity peaks during the morning and evening, although the timing of the evening peak varied by photoperiod (Fig. 1c). We quantified for each fly the average evening peak time of the last 4 of 7 days in a given photoperiod, allowing the first 3 days to serve as an acclimation period (Fig. 1c). For both *D. melanogaster* strains, we observed that as photoperiod increases, the timing of the evening activity peak is commensurately delayed (Fig. 1c). By contrast, for our *D. sechellia* strains, we observed strikingly little photoperiod plasticity, with a median delay in evening peak time of maximum ∼1 h regardless of photoperiod length (Fig. 1c). Additionally, under all photoperiod regimes, *D. sechellia* ended its afternoon siesta and began ramping up to evening peak activity a few hours earlier than *D. melanogaster* (Fig. 1c).

### *D. sechellia* displays reduced morning peak activity compared to *D. melanogaster*

We noted that *D. sechellia* is far less active during the dark phase than either *D. melanogaster* strain and displays little, if any, morning anticipation, i.e., increasing activity in the hours leading up to lights-on (Fig. 1c-d). Quantification of pre-dawn activity under 12:12 h LD in the 3 h preceding lights-on revealed prominent differences between the species (Fig. 1d): *D. sechellia* is generally very inactive during this time period, while the *D. melanogaster* strains display ample, albeit strain-specific, activity. These observations led us to question whether the morning peak of *D. sechellia* is a true activity peak or merely a startle response to lights-on. To address this issue, we measured free-running activity by acclimating our strains to 12:12 h LD before submitting them to constant dark conditions (DD). Both *D. melanogaster* and *D. sechellia* remained rhythmic under DD (Fig. 1e), and each strain displayed a period of ∼24 h (Extended Data Fig. 1). By contrast, while *D. melanogaster* displayed clear activity peaks at the subjective dawn, *D. sechellia* exhibited very little activity at this timepoint, even during the first day of DD (Fig. 1e). This result supports the hypothesis that the morning peak of *D. sechellia* observed under LD conditions (Fig. 1c) is predominantly a startle response to lights-on. Consistently, when we quantify morning peak timing under 12:12 h LD, we found that *D. melanogaster* reached peak activity before lights-on, as previously described^17^, while *D. sechellia* peaked only at or after lights-on (Fig. 1f).

### Reduced circadian plasticity and morning activity represent an evolutionary loss in *D. sechellia*

To determine if the species differences in circadian plasticity and morning activity represent an evolutionary loss in *D. sechellia* or a trait gain in *D. melanogaster*, we measured the activity of additional *D. melanogaster* strains collected from the Lower Zambezi Valley (close to its ancestral range^18^ (∼16° S), as well as strains of two species that have a more recent common ancestor with *D. sechellia* (Fig. 1a): *D. simulans* (collected from Madagascar, its ancestral range^19^, ∼19° S) and *D. mauritiana* (a species endemic to Mauritius, ∼20° S) (Extended Data Fig. 2a). Comparing 12:12 h and 16:8 h LD conditions, we observed a similar larger degree of circadian plasticity (∼2 h evening peak delay under the longer photoperiod) for all of these strains compared to *D. sechellia* (∼1 h evening peak delay). All of these strains also exhibited significant morning anticipation (Extended Data Fig. 2b). These results indicate that the lack of plasticity and reduction in morning peak activity observed in *D. sechellia* likely represent evolutionary losses in this lineage, and point to a potential connection between these two phenotypes.

**Fig. 2.**
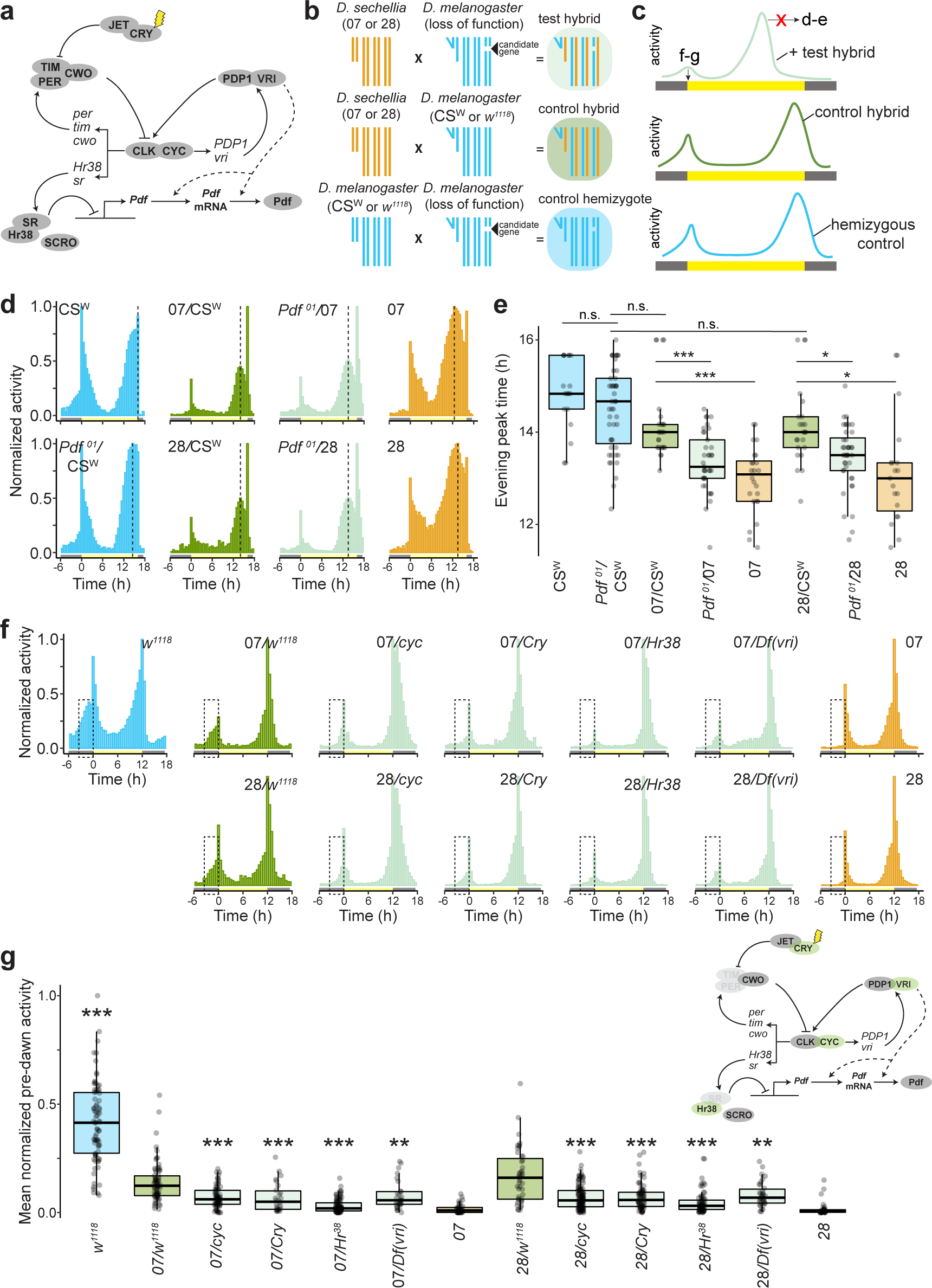
A screen of circadian clock genes reveals distinct genetic architectures underlying interspecific differences in plasticity and morning activity. **a**, Molecular components of the circadian clock in *D. melanogaster*. **b**, Crossing schemes used to generate hemizygous test hybrids, heterozygous control hybrids, and hemizygous *D. melanogaster* flies in a controlled genetic background. The fourth (“dot”) chromosome is not shown. **c**, Schematics illustrating the sought-after behavioural phenotypes of test hybrids, and anticipated phenotypes of control hybrids and hemizygous *D. melanogaster* controls: positive candidate gene test hybrids, but not corresponding controls, will display reduced circadian plasticity and/or reduced morning activity. **d**, Mean normalized activity of the indicated control and hybrid genotypes under a 16:8 h LD cycle. Plots depict average activity of the last 4 days of a 7-day extended photoperiod, following 7 days of 12:12 h LD. Vertical dashed lines indicate the average timing of the evening peak for each strain. Sample sizes as follows: CS^W^ (16), *Pdf^01^*/CS^W^ (47), 07/CS^W^ (25), 28/CS^W^ (23), 07/*Pdf^01^* (37), 28/*Pdf^01^*(40), 07 (24), 28 (19). Full screen results are shown in Extended Data Fig. 3. **e**, Evening peak time for the flies depicted in **d**. Asterisks indicate significant differences: * = p < 0.05 and *** = p < 0.001 (Wilcoxon tests with Bonferroni correction). Comparisons were made only between control and test hybrids of the same genetic backgrounds. **f**, Mean normalized activity of the indicated genotypes under a 12:12 h LD cycle, illustrating the screened mutations displaying reduced morning anticipation in test hybrids. Dashed boxes highlight the pre-dawn area used to quantify morning anticipation. Full screen results are shown in Extended Data Fig. 4. **g**, Mean normalized pre-dawn activity for the genotypes in **f.** Asterisks indicate significant differences: ** = p < 0.01 and *** = p < 0.001 (Wilcoxon tests comparing each test hybrid to the control hybrid strain (07/*w^1118^*) with Bonferroni correction). Top right: the circadian molecular network in which screen hits for morning anticipating are highlighted in green; genes in light grey were unable to be tested (see Methods).

### A genetic screen for differences in circadian activity and plasticity

To identify the mechanistic basis of the species differences in circadian behaviors, we employed a candidate genetic screening approach. Extensive research in *D. melanogaster* has defined central brain circuitry of 150 circadian neurons, divided into discrete groups with differing effects on circadian activity and network dynamics^20^. Within each neuron, a gene regulatory feedback loop allows each cell to track a ∼24 h period^21-23^ and to control the rhythmic expression of downstream effector genes (Fig. 2a). The members of this network serve as excellent candidates for explaining species-specific circadian behaviors. To take as unbiased an approach as possible, we obtained loss-of-function mutations for the majority of the genes encoding proteins within this feedback loop in addition to several in the downstream network, including the neuropeptide Pigment-dispersing factor (Pdf), as well as other circadian neuropeptides, CCHa1 and ITP (Supplementary Table 1). Our cross-species behavioral analyses (Fig. 1c-d and Extended Data Fig. 2) indicated that reduced circadian plasticity and morning peak activity in *D. sechellia* represent evolutionary losses. We therefore reasoned that the causal *D. sechellia* alleles were more likely to be recessive to *D. melanogaster*, and designed a screen in *D. melanogaster-D. sechellia* hybrids (Fig. 2b). In brief, we generated hemizygous test hybrids containing *D. melanogaster* mutants for each individual candidate gene, to reveal the recessive phenotype of the *D. sechellia* allele at the same locus. We also generated heterozygous control hybrids, using the *D. melanogaster w^1118^* strain (the most common genetic background of our mutant strains) or CS^W^ strain (in the case of *Pdf*; see Methods) and each of our *D. sechellia* strains. These control hybrids have one allele each from *D. melanogaster* and *D. sechellia*. Thus, any differences we observe between control and test hybrids is likely due to the loss of the *D. melanogaster* allele in the test hybrid background. To control for genetic background effects^24^, we tested hybrids of both the *Dsec*07 and *Dsec*28 backgrounds. Finally, gene dosage effects were assessed by testing control hemizygotes in a (non-hybrid) *D. melanogaster* background, i.e., mutants crossed to *w^1118^* or CS^W^. Genes whose mutations displayed an effect in both test hybrid backgrounds compared to control hybrids, and no effect in hemizygous *D. melanogaster*, were considered the strongest candidates explaining interspecific phenotypic differences (Fig. 2c).

### The *Pigment-dispersing factor* gene underlies evolved differences in circadian plasticity

To assess candidate genes for an effect on circadian plasticity, we observed test and control hybrids under a 16:8 h LD photoperiod. Control hybrids of either the *w^1118^* (Extended Data Fig. 3a-d) or CS^W^ background (Fig. 2d-e) display a larger degree of phenotypic plasticity than their *D. sechellia* parental strain, confirming that the *D. melanogaster* genotype underlying plasticity is at least partially dominant to that of *D. sechellia*, though the degree of dominance depends on the *D. melanogaster* parental strain. We screened 14 genes covering the majority of the circadian transcriptional feedback loop and many of its modulator and effector genes. Mutations in only one reduced circadian plasticity in both test hybrid backgrounds but not in hemizygous *D. melanogaster*: *Pdf* (Fig. 2d-e, Extended Data Fig. 3e-f). This is a promising gene for explaining species differences because, in *D. melanogaster*, Pdf is essential for delaying the phase of the endogenous clock circadian neurons under long photoperiods^25^, and flies lacking Pdf expression display reduced plasticity^26,27^.

**Fig. 3.**
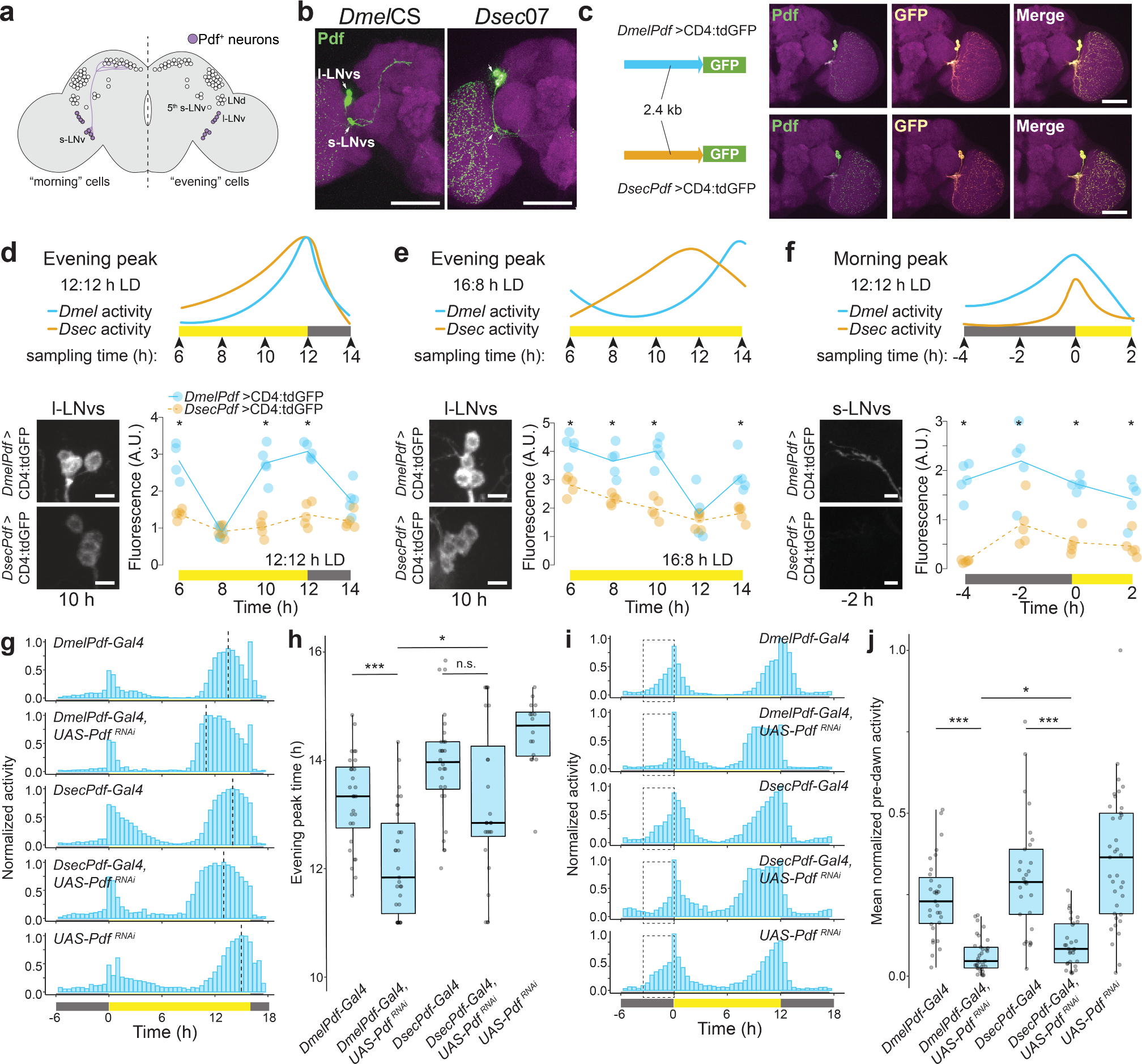
Species-specific *cis*-regulatory elements impact *Pdf* expression and have the potential to impact behaviour. **a**, Schematic of the circadian clock network in *D. melanogaster*, which is composed of ∼75 neurons in each brain hemisphere that are divided into distinct groups. The groups comprising the morning and evening cells are indicated on the left and right hemispheres, respectively. Pdf-positive small and large ventrolateral neurons (s-LNvs and l-LNvs) are highlighted in purple. **b**, Immunofluorescence for Pdf (green) and Cadherin-N (magenta) on whole-mount brains of the indicated strains at 2 h under 12:12 h LD conditions. **c**, Representative images of reporter expression visualized by Pdf (left) and GFP immunofluorescence (middle) showing faithful labelling (merge, right) of s-LNvs and l-LNvs for both the *D. melanogaster* (top) and *D. sechellia* (bottom) *Pdf* 5’-regulatory regions. For **b** and **c**, scale bars, 100 μm. **d**, Top: Schematic illustrating the average activity patterns of *D. melanogaster* and *D. sechellia* during behaviourally relevant time points (labelled with arrowheads) within the evening peaks under 12:12 h LD conditions where we analyzed *Pdf* expression. These summaries were derived from the data in Fig. 1c-d. Bottom-left: representative images of GFP immunofluorescence in the l-LNvs for the *D. melanogaster* and *D. sechellia Pdf* 5’-regulatory sequence-GFP reporter strains under 12:12 h LD at one time point (10 h). Bottom-right: GFP fluorescence quantifications at 5 time points spanning the evening activity peak period. **e**, Top: Schematic illustrating the average activity patterns of *D. melanogaster* and *D. sechellia* during behaviourally relevant time points (labelled with arrowheads) within the evening peaks under 16:8 h LD conditions where we analyzed *Pdf* expression. Bottom-left: representative images of GFP reporter immunofluorescence in the l-LNvs for the *D. melanogaster* and *D. sechellia* strains under 16:8 h LD at one time point (10 h). Bottom-right: GFP fluorescence quantifications at 5 time points spanning the evening activity peak period. **f**, Top: Schematic illustrating the average activity patterns of *D. melanogaster* and *D. sechellia* during behaviourally relevant time points (labelled with arrowheads) within the morning peaks under 12:12 h LD conditions where we analyzed *Pdf* expression. Bottom-left: representative images of GFP immunofluorescence in the s-LNv axon terminals for the *D. melanogaster* and *D. sechellia Pdf* reporter strains at one time point (-2 h). We observed the same general pattern when measuring fluorescence in the s-LNv soma. Bottom-right: GFP fluorescence quantifications at 4 time points spanning the morning activity peak period. For **d**-**f**, N = 5 for each strain and time point. Despite weak signal in some *D. sechellia* images, the projections were easily identified in threshholded images. **g**, Mean normalized activity of the indicated *D. melanogaster* genotypes under a 16:8 h LD cycle. Plots depict average activity of the last 4 days of a 7-day extended photoperiod, following 7 days of 12:12 h LD. Vertical dashed lines indicate the average timing of the evening peak for each strain. Sample sizes as follows: *DmelPdf-Gal4/+* (28), *DsecPdf-Gal4/+* (28), *UAS-Pdf^RNAi^/+* (16), *DmelPdf-Gal4/UAS-Pdf^RNAi^* (31), *DsecPdf-Gal4/UAS-Pdf^RNAi^* (24). **h**, Evening peak time for the flies shown in **g**. **i**, Mean normalized activity of the indicated *D. melanogaster* genotypes under a 12:12 h LD cycle. Plots depict average activity of the last 4 days of a 7-day recording period. Dashed boxes highlight the pre-dawn period, 3 h before lights-on. Sample sizes as follows: *DmelPdf-Gal4/+* (31), *DsecPdf-Gal4/+* (29), UAS-*Pdf^RNAi^*/+ (47), *DmelPdf-Gal4/UAS-Pdf^RNAi^* (34), *DsecPdf-Gal4/UAS-Pdf^RNAi^* (30). **j**, Mean normalized pre-dawn activity for the genotypes in **i**. For **d**-**f**, Lines connect medians of each time point within genotypes. Scale bars, 10 μm. For **d**-**f, h** and **j**, Asterisks indicate significant differences: * = p < 0.05 and *** = p < 0.001 (Wilcoxon tests with Bonferroni correction).

### Potential broad-scale divergence of the circadian clock underlies *D. sechellia*’s reduced morning peak activity

We also screened these genotypes under 12:12 h LD and quantified pre-dawn activity (Extended Data Fig. 4a-d). In contrast to the dominance of the *D. melanogaster* phenotype for circadian plasticity (Extended Data Fig. 3a-d), *w^1118^* control hybrids display intermediate pre-dawn activity relative to either parental strain, suggesting a more complex genetic architecture of this species difference. Consistent with this idea, four genes displayed an effect in test hybrids of both backgrounds (Fig. 2f-g) and not in hemizygous *D. melanogaster* flies (Extended Data Fig. 4e-g). These encode the core transcriptional feedback loop protein CYC and the light-sensitive CRY, which is responsible for light-dependent synchronization of the molecular clock^28-30^, as well as Hr38 and VRI, which are neural activity-dependent transcriptional or post-transcriptional regulators of *Pdf* expression, respectively^31^.

**Fig. 4.**
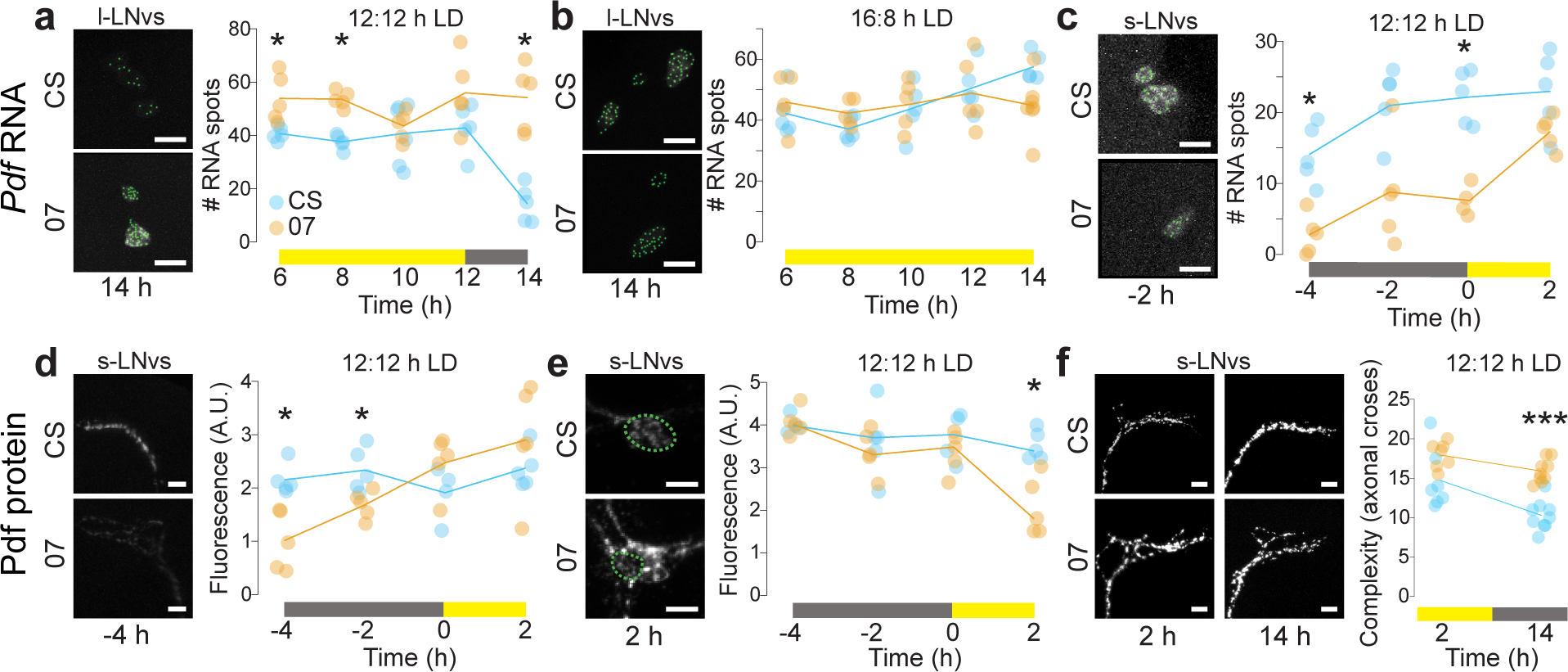
*Pdf* expression differences observed between *D. melanogaster* and *D. sechellia*. **a**, Left: representative images of *Pdf* smFISH in the l-LNv soma in the CS and 07 strains under 12:12 h LD at one time point (14 h), with RNA spots (green) identified by RS-FISH. Right: quantifications of RNA spots at 5 time points spanning the evening activity peak period. **b**, Left: representative images of *Pdf* smFISH in the l-LNv soma in the CS and 07 strains under 16:8 h LD at one time point (14 h), with RNA spots (green) identified by RS-FISH. Left: quantifications of RNA spots in the CS and 07 strains under 16:8 h LD at 5 time points spanning the evening activity peak period. **c**, Left: representative images of smFISH in the s-LNv soma for the CS and 07 strains under 12:12 h LD at one time point (-2 h) with RNA spots identified by RS-FISH. Right: quantifications of RNA spots in the 07 and CS strains at 4 time points spanning the pre-dawn period. **d**, Left: representative images of Pdf immunofluorescence in the s-LNv axon terminals for the CS and 07 strains at one time point (-4 h). Right: quantifications of Pdf signals at 4 time points spanning the morning activity peak period. **e**, Left: representative images of Pdf immunofluorescence in the s-LNv cell bodies for the CS and 07 strains at one time point (2 h). Right: quantifications of Pdf signals at 4 time points spanning the morning activity peak period. **f**, Left: representative images of Pdf immunofluorescence in the s-LNv axon terminals for the CS and 07 strains during the day (2 h) and night (14 h). Right: quantifications of axonal branching complexity quantifications. For **a**-**f**, N = 5 brains per strain per time point. Plotted values are the average of left and right hemispheres. Lines connect medians of each time point within genotypes. Asterisks indicate significant differences: * = p < 0.05 and *** = p < 0.001 (Wilcoxon tests with Bonferroni correction). All scale bars, 10 μm.

### *Cis*-regulatory evolution of *Pdf*

We subsequently focused our attention on *Pdf*, because of its unique significant effect on circadian plasticity and evidence that *trans*-regulation of *Pdf* expression influences morning peak activity. To understand how this gene differs between species, we first compared the *Pdf* coding sequence of 10 *D. melanogaster* and 6 *D. sechellia* isogenic strains (as well as 5 *D. simulans* lines). These sequences are predicted to encode peptides of near-perfect conservation, with no species-specific differences (Extended Data Fig. 5).

**Fig. 5.**
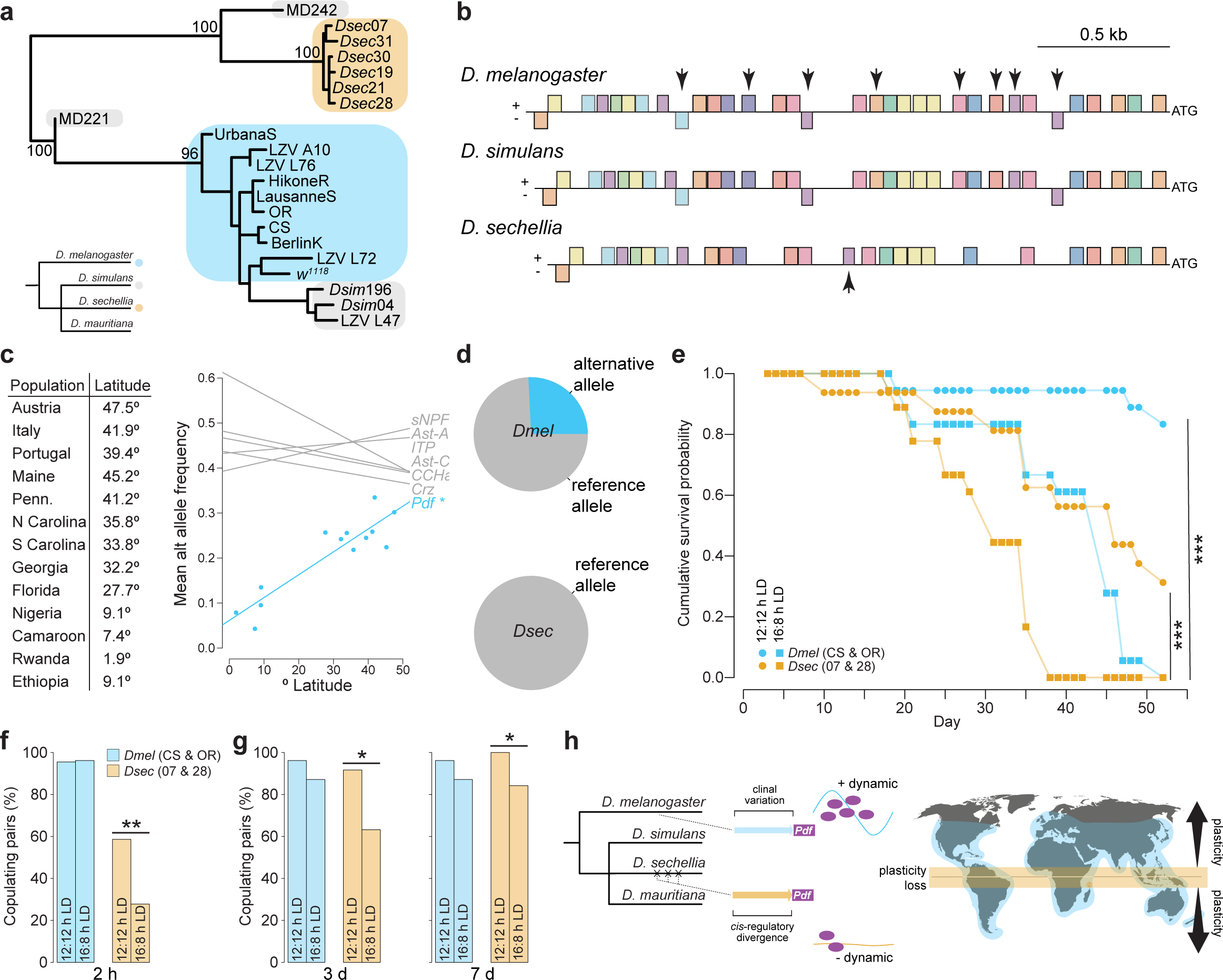
Evidence for selection on the *Pdf* 5’-regulatory sequence and fitness effects of circadian plasticity loss. **a**, A midpoint rooted maximum likelihood phylogeny of *Pdf* 5’-regulatory sequences from 10 *D. melanogaster* (blue), 6 *D. sechellia* (orange), and 5 *D. simulans* (grey) strains. Bootstrap support values are shown for key internal nodes (100 bootstraps). The species tree is depicted at the bottom left for comparison. **b**, A motif analysis of the *Pdf* 5’-regulatory sequences from **a**. The *Pdf* start codon is on the right, and the different types of predicted regulatory motifs for each species are shown as distinct colored boxes on the + or – strand. Species-specific diagrams depict all motifs found for each species. Motifs observed in *D. melanogaster* and *D. simulans* but absent in all *D. sechellia* sequences are marked with downward facing arrows; one motif unique to all sequences of *D. sechellia* is marked with an upward facing arrow. No variation in motif location was observed among the 6 *D. sechellia* strains. **c**, Left: the 13 *D. melanogaster* populations selected from Ref.^48^ and the approximate latitude of their collection sites. Right: plot of minor allele frequency in the *Pdf* 5’-regulatory sequences (blue) of these 13 populations against latitude, revealing a significant positive correlation (Spearman’s rho = 0.77). No such correlation is observed for the putative 5’-regulatory sequences of 6 control neuropeptide genes (grey); detailed data points are shown in Extended Data Fig. 8. **d**, Average minor allele frequency of variable sites from the analysis in **c** in the laboratory *D. sechellia* and *D. melanogaster* lines from **a**. Variable sites are significantly underrepresented in *D. sechellia* relative to *D. melanogaster* strains (p <0.05, Fisher’s exact test). **e**, Cumulative survival probability for *D. melanogaster* (*Dmel*CS + *Dmel*OR, blue) and *D. sechellia* (*Dsec*07 + *Dsec* 28, orange) maintained at 12:12 h LD (circles) or 16:8 h LD (squares) for 52 days. No significant differences were observed between strains of the same species by photoperiod, and were thus pooled (Fisher’s exact test, all p > 0.05). Pooled data were compared between photoperiods within species using a log-rank test. **f**, Percent of copulating pairs observed for *D. melanogaster* (*Dmel*CS + *Dmel*OR, blue) and *D. sechellia* (07 + 28, orange) after 2 h for flies acclimated to 12:12 h LD (left) compared to 16:8 h LD (right, Wilcoxon test). No significant differences were observed between strains of the same species by photoperiod, and were thus pooled (Fisher’s exact test, all p = 1). Sample sizes as follows: *D. melanogaster* 12:12 h LD (22), *D. melanogaster* 16:8 h LD (26), *D. sechellia* 12:12 h LD (29), and *D. sechellia* 16:8 h LD (32). **g**, Same as **f**, except after 3 days (left) or 7 days (right). Sample sizes as follows: *D. melanogaster* 12:12 h LD (26), *D. melanogaster* 16:8 h LD (31), *D. sechellia* 12:12 h LD (36), and *D. sechellia* 16:8 h LD (38). **h**, Schematic of the main findings of this work: the equatorial species *D. sechellia* has lost circadian plasticity, in part through *cis*-regulatory changes in the *Pdf* 5’ region, which lead to less dynamic expression. For **c, e**-**g**, Asterisks indicate significant differences: * = p < 0.05, ** = p < 0.01, and *** = p < 0.001.

We therefore reasoned that divergence between species must be due to expression differences of Pdf. In *D. melanogaster*, this neuropeptide is expressed exclusively in 8 neurons in each central brain hemisphere: 4 large ventrolateral clock neurons (l-LNvs), which represent a subset of the “evening” cells, and 4 small ventrolateral clock neurons (s-LNvs), the “morning” cells (Fig. 3a). These neuronal subtypes have predominant roles in controlling evening and morning activity, respectively^32-34^, although a functional clock is required in both for photoperiod plasticity^33,35^. Through qualitative Pdf immunofluorescence analysis, we observed a conserved spatial pattern of Pdf expression in *D. sechellia* (Fig. 3b), consistent with a survey of Pdf expression across a broader range of drosophilids^36^. This result suggested that species-specific differences instead exist in the temporal pattern and/or levels of expression.

Because our hybrid screen identified an effect of the *Pdf* locus itself, we hypothesized that differences in expression must result from divergence in the *cis*-regulatory region. To test this possibility, we cloned ∼2.4 kb of genomic DNA immediately 5’ of the start codon of *Pdf* from either *D. melanogaster* or *D. sechellia* – based upon a previous analysis in *D. melanogaster^37^* – upstream of a GFP reporter gene^38^. These transgenes were integrated in an identical genomic location in the same *D. melanogaster* genetic background, thereby permitting comparison of their activity in a common *trans* and genomic environment. As expected, both species’ *Pdf* reporters exclusively labelled the l-LNvs and s-LNvs (Fig. 3c). We first measured reporter expression in the l-LNvs because, in *D. melanogaster*, Pdf expression in these cells is required to plasticly adjust the timing of the evening peak^4^. To process all samples in parallel for quantitative comparisons, we focused on behaviorally relevant time points under 12:12 h LD and 16:8 h LD conditions (Fig. 3d-e). In the l-LNvs, throughout the evening activity peak under both photoperiods, the *D. sechellia* 5’-regulatory region consistently drives lower and more constant reporter expression relative to the *D. melanogaster* sequence (Fig. 3d-e). Notably, the *D. melanogaster Pdf* reporter displayed a sudden drop in expression at 8 h (under 12:12 h LD) or at 12 h (under 16:8 h LD), prior to returning to a higher level, which potentially reflects a new pulse in transcriptional activity that appears sensitive to photoperiod.

In *D. melanogaster*, the s-LNvs are essential for resetting the phase of the circadian clock (likely through the cyclic release of Pdf^39,40^) and are necessary for morning peak activity^32,34^. We therefore compared reporter expression in the s-LNv axonal projections – where the largest cyclic Pdf expression is observed over a 24 h period^37^ – for four time points spanning the morning activity peak (Fig. 3f). We again observed that the *D. sechellia* sequence drives lower expression of the reporter but, in contrast to the l-LNvs during the evening peak, with a similar temporal fluctuation in expression. Together these results confirm functional divergence of the 5’ *cis*-regulatory region between *D. sechellia* and *D. melanogaster*. This sequence is most likely to affect transcriptional activity but, because it encompasses the 5’-UTR of *Pdf*, we cannot exclude that it (also) influences transcript stability and/or translatability^41^.

### Modifying *Pdf* expression reduces the magnitude of circadian plasticity and morning anticipation

Having identified species-specific properties in the *cis*-regulatory element of *Pdf*, we tested whether these would be sufficient to impact behavior. Taking advantage of the different reporter expression levels driven by the *D. melanogaster* and *D. sechellia Pdf* 5’ regions (Fig. 3d-f), we used these same sequences to generate Pdf neuron Gal4 drivers to induce “strong” (*D. melanogaster Pdf-Gal4*) or “weak” (*D. sechellia Pdf-Gal4*) knock-down of *Pdf* using a *UAS-Pdf^RNAi^* effector^42^. We first validated the anticipated distinct efficacy of RNAi with quantitative single molecule RNA fluorescence *in situ* hybridization (smFISH). In *DmelPdf*-Gal4>*Pdf* RNAi flies, *Pdf* transcripts were reduced to ∼4% of the levels of control animals, while *DsecPdf*-Gal4>*Pdf* RNAi flies expressed *Pdf* transcripts at ∼20% of control levels (Extended Data Fig. 6).

We first observed these flies under 16:8 h LD conditions and quantified evening peak timing (Fig. 3g-h). The *DmelPdf*-Gal4>*Pdf* RNAi flies displayed a dramatic reduction in evening peak time relative to controls. By contrast, *DsecPdf*-Gal4>*Pdf* RNAi flies display a small, and non-significant, decrease in evening peak time relative to control animals, with a notable increase in variance. Importantly, there is a significant difference between the RNAi-expressing genotypes despite their otherwise identical genetic backgrounds. We also observed the behavior of these flies under 12:12 h LD, and quantified pre-dawn activity (Fig. 3i-j). Both *DmelPdf*-Gal4>*Pdf* RNAi and *DsecPdf*-Gal4>*Pdf* RNAi flies have reduced pre-dawn activity relative to controls. However, the *DmelPdf*-Gal4>*Pdf* RNAi flies displayed significantly less pre-dawn activity than the *DsecPdf*-Gal4>*Pdf* RNAi flies. Together these results indicate that the level (and possibly temporal dynamics) of *Pdf* expression – as determined by species-specific 5’ *cis*-regulatory regions – is sufficient to affect both circadian plasticity and morning anticipation in an otherwise identical genetic background.

### Species-specific, photoperiod-dependent differences in *Pdf* RNA expression

We next investigated how *cis*-regulatory divergence might influence endogenous *Pdf* RNA and protein expression. Using smFISH, we compared *Pdf* transcript levels between *D. melanogaster* and *D. sechellia* at the same fine temporal resolution as for the transgenic reporters. Under 12:12 h LD, quantification of transcript levels in l-LNvs throughout the evening peak revealed slightly higher initial *Pdf* expression in *D. sechellia* than *D. melanogaster* (Fig. 4a). The most striking difference, however, is a sudden drop in *Pdf* transcripts in *D. melanogaster*, but not *D. sechellia*, after lights-off (Fig. 4a). Under 16:8 h LD conditions, this reduction in *Pdf* RNA levels is no longer present (Fig. 4b). These observations indicate that transcriptional activity of *Pdf* is more dynamic in *D. melanogaster* than *D. sechellia*, resulting in a decrease of *Pdf* transcripts by the dark phase under 12:12 h LD. Such a pattern might additionally/alternatively result from differences in RNA stability between species. Regardless of the mechanism, this species-specific transcript depletion appears sensitive to photoperiod. Interestingly, such a pattern of *Pdf* transcription in the l-LNvs has not been previously described using bulk or single-cell RNA sequencing of these neurons in *D. melanogaster*^43,44^ when sampling more broadly across a 24 h time period; however a decrease in *Pdf* RNA at 14 h (relative to the 2 h time point) was previously documented using smFISH^45^, congruent with our results.

When we quantified *Pdf* RNA in the s-LNvs throughout the morning activity peak, we found that *Dmel*CS displayed overall more *Pdf* RNA than *Dsec*07 at each time point, particularly in the pre-dawn time points, with *Dsec*07 reaching near-similar levels only after lights-on (Fig. 4c). This difference in *Pdf* RNA levels is concordant with the differences in pre-dawn activity we observed between these species (Fig. 1c-d), and the implication of transcriptional and post-transcriptional regulators of *Pdf* expression in species-specific morning anticipation (Fig. 2f-g).

### Species- and photoperiod-dependent differences in Pdf protein expression

We next used immunofluorescence to compare Pdf protein expression in *D. melanogaster* and *D. sechellia*. Similar to the transcript analyses, we quantified Pdf immunofluorescence in the l-LNvs in time points surrounding the evening activity peak under 12:12 h LD and 16:8 h LD conditions (Extended Data Fig. 7a-b). Under both photoperiods, we observed qualitative differences between these species in the overall pattern of staining intensity, but the much greater variability in signal intensity of these samples, particularly for *D. sechellia* under 16:8 h LD, made it difficult to connect back to our more quantitative measures of *Pdf* RNA levels (or to behavioral activity).

In the s-LNvs, we quantified Pdf fluorescence in the axonal projections for the same time points spanning the morning activity peak (Fig. 4d). In the relatively short time window analyzed, we observed a consistently high level of Pdf in *D. melanogaster*, including in the hours preceding lights-on. By contrast, in *D. sechellia*, Pdf signal is lower in the pre-dawn hours, and increases to an equivalent amount as *D. melanogaster* only by lights-on. This pattern corresponds well to that of the relative levels of *Pdf* transcripts, and to species differences in pre-dawn activity at these times (Fig. 1c-d). We also analyzed Pdf immunofluorescence in the s-LNv cell bodies, which display higher and less cyclic Pdf expression in *D. melanogaster*^37^. Consistently, we found Pdf signal remains high across the morning peak times in the s-LNv soma of *D. melanogaster* (Fig. 4e). In *D. sechellia*, the Pdf signal begins high and drops significantly only after lights-on (Fig. 4e). Together, our observations of *Pdf* RNA levels and Pdf protein levels/distribution in the s-LNvs of *D. sechellia* suggest that this species has a smaller and/or shorter pulse of *Pdf* expression around the morning peak compared to *D. melanogaster*, such that once this neuropeptide accumulates to high levels in the axon terminals at/after lights-on (Fig. 4d), it becomes depleted in the soma (Fig. 4e). Species-specific dynamics in Pdf spatial distribution might also reflect differences in intracellular transport and/or secretion pathways.

### Species-specific circadian structural plasticity of Pdf neurons

In *D. melanogaster*, the axonal projections of the s-LNvs to the dorsal circadian neurons (Fig. 3a) display circadian structural plasticity, reaching peak branching complexity during the day, and lower complexity during the night^46^. This phenomenon depends, at least in part, on cyclic expression and release of Pdf from both the s-LNvs and l-LNvs and the expression of the Pdf receptor^42^. To test if the observed species-specific temporal patterns of Pdf expression are accompanied by differences in the remodeling of these neurons, we quantified the branching complexity of s-LNv projections in *D. melanogaster* and *D. sechellia* during the light (2 h) and dark (14 h) phases (Fig. 4f). During the light phase, we found statistically indistinguishable levels of complexity in the two species. However, in *D. melanogaster*, branching complexity is significantly lower in the dark phase when compared to *D. sechellia* (Fig. 4f). This apparent reduction of structural plasticity of *D. sechellia* Pdf neurons corroborates the less dynamic changes in Pdf expression in this species.

### Signals of *cis*-regulatory evolution and selection on the *Pdf* regulatory region

Having characterized functional divergence of the *Pdf cis*-regulatory region between species, we next asked whether the *D. sechellia* sequence exhibits any evolutionary signature. We sequenced this region in all of our *D. melanogaster*, *D. sechellia* and *D. simulans* strains. Overall, *D. simulans* and *D. melanogaster* sequences share an average of ∼97% pairwise sequence similarity, while *D. sechellia* has an average pairwise sequence similarity of ∼93% with *D. melanogaster*. We used these sequences to construct a maximum likelihood phylogeny (Fig. 5a). In contrast to the species tree, the *Pdf* 5’-regulatory sequences from *D. sechellia* form a monophyletic group, while the *D. melanogaster* and *D. simulans* sequences mostly cluster together (with the exception of a single *D. simulans* sequence, which groups more basally with the *D. sechellia* sequences). Motif enrichment analysis^47^ identified putative regulatory sequences in these species’ 5’ regions. While all such motifs were shared among the *D. melanogaster* and *D. simulans* sequences, 8 of these sites are degenerated or absent in *D. sechellia* (and one site is unique to this species) (Fig. 5b), indicating that sequence divergence in the *D. sechellia Pdf* regulatory region is likely to affect its function activity, potentially through the loss of transcription factor binding sites. We next investigated whether the sequence divergence between *D. melanogaster* and *D. sechellia* 5’-regulatory sequences might result, at least in part, from natural selection on variants underlying circadian plasticity at higher latitudes. We examined this possibility by determining whether variants within the *D. melanogaster Pdf* 5’-regulatory region are associated with higher degrees of circadian plasticity observed with increasing latitudes^15^. The reverse analysis in *D. sechellia* is not possible as this species is restricted to an equatorial latitude. Taking advantage of a dataset of single nucleotide variant frequencies in the genomes of globally-distributed populations of *D. melanogaster*^48^, we chose 13 populations representing a wide range of latitudes (Fig. 5c); all had a minimum read-depth of 5 to ensure confidence in variant frequencies. We calculated the average minor allele frequency (MAF) across all variable sites detected within the *Pdf* 5’-regulatory sequence and plotted these against the estimated latitude of the population collection site. Because correlations could reflect the underlying population structure as a result of demographic history as *D. melanogaster* emigrated from its native range in Africa^49^, as a control, we repeated this analysis for the putative equivalent regulatory region (2.4 kb upstream from the start codon) of six other neuropeptide genes. We found a strong positive correlation (Spearman’s rho = 0.77) between population latitude and MAF for the *Pdf* 5’-regulatory region, but not for any of the other neuropeptide genes (Fig. 5c, Extended Data Fig. 8). These comparisons indicate that the effect of latitude on MAF of the *Pdf* 5’-regulatory sequence is different than we would expect due to demography alone, suggesting a potential role for selection on these sites in *D. melanogaster*. These results are similar to recent reports of clinal variation in other circadian genes^50-53^.

Lastly, we checked the MAFs of these variable sites in our laboratory strains. These single nucleotide variants occur with a MAF of ∼25% among our *D. melanogaster* strains, but none were present in any of our *D. sechellia* strains (Fig. 5d). These results are consistent with a potential function of these variants in increasing circadian plasticity. Together with the predicted motif differences between *D. melanogaster* and *D. sechellia Pdf* 5’-regulatory sequences (Fig. 5b), these data will help guide future analyses of the specific functional changes within this region.

### Circadian plasticity is important for reproductive success

To identify a mechanism by which natural selection might act, we asked if plasticity in circadian activity impacts fitness. Photoperiod has been shown to affect lifespan in many insects and other animals^54-56^, leading us to reason that if flies have reduced lifespans under extended photoperiod, then this might lead to reduced total reproductive output. We recorded survivorship of individual *D. melanogaster* and *D. sechellia* maintained at either 12:12 h or 16:8 h LD (Fig. 5e). Flies of both species maintained under 16:8 h LD displayed a significant reduction in lifespan relative to those under 12:12 h LD. This result is surprising in that it suggests that *D. melanogaster*’s ability to plasticly adjust its activity to longer days does not alleviate the cost of exposure to longer photoperiod. However, the detrimental effect of longer photoperiod was not observed until after several weeks in both species, in which time flies could certainly mate and produce offspring. It is therefore unclear if the effect of extended photoperiod on lifespan would impact fitness in nature, where lifespan is likely much shorter than under laboratory conditions.

Circadian rhythms are important for synchronizing sexual behavior among conspecifics^10,11^. We therefore reasoned that if circadian plasticity (or the lack thereof) impacted copulation success, it would likely impact fitness. To test this possibility, we acclimated male and female *D. melanogaster* and *D. sechellia* virgins to 12:12 h LD and 16:8 h LD for 4 days. We then observed copulation rates among male-female pairs in two assays: first, over a 2-hour period just after lights-on (Fig. 5f), and second, over the course of one week (Fig. 5g). For *D. melanogaster*, we observed no difference in copulation rates of flies between treatments in either experiment. By contrast, there was a significant decrease in copulation by *D. sechellia* acclimated to 16:8 h LD. Specifically, the decrease in copulation rate in the short-term assay was largely maintained over the course of several days in our long-term assay, indicating that flies that did not copulate within 2 h were unlikely to do so several days later. These results demonstrate that *D. sechellia*’s reproductive output – and thus fitness – is highly likely to be impacted by its lack of circadian plasticity under extended photoperiods that it would never normally experience in nature. These data also suggest that by plasticly adjusting its behavior, *D. melanogaster* is able to circumvent these negative effects.

## Discussion

Identifying the genetic and neural mechanisms of behavioral plasticity is key to understanding how organisms evolve(d) to inhabit variable environments, as well as to projecting how they will persist in increasingly unstable ones^57^. However, efforts to uncover the proximate causes of behavioral divergence are limited by a lack of genetic access to multiple closely-related species, leaving remarkably few cases where the molecular and/or cellular underpinnings of interspecific differences in behaviors have been mapped^58^, with the vast majority being in (peripheral) sensory pathways (e.g.,^59-63^). Here, we have uncovered molecular and cellular mechanisms of circadian plasticity differences in drosophilids, providing a rare example linking differences in gene function, central neuron populations, and behavioral differences between species.

By comparing the equatorial *D. sechellia* with closely-related, globally-distributed species, we discovered a dramatic difference in the degree of photoperiod plasticity and provided evidence for a key contribution of the *cis*-regulatory region of the *Pdf* locus in this difference (Fig. 5h). In *D. melanogaster*, Pdf expression is required in the l-LNvs for photoperiod plasticity^27^, and previous comparative work has described interspecific spatial differences in Pdf expression^36^, notably in high-latitude species where Pdf is restricted to the l-LNvs^4,16^. Importantly, prior to our work, no functional connection between divergence at the *Pdf* locus and species differences in behavior had been established. Nevertheless, these observations, combined with our analyses in *D. sechellia* and *D. melanogaster* – including the evidence for latitude-based selection on the *D. melanogaster Pdf* 5’-regulatory region – point to the *Pdf* locus as a hotspot of evolution. Given Pdf’s terminal placement as an effector gene of the clock network^31^, its role in broadly synchronizing circadian clock neurons^64^, and its strong impact on circadian behaviors, changes in the *cis*-regulation of *Pdf* expression might represent a minimally pleiotropic means of introducing plasticity into the clock neuronal network, akin to regulatory changes of developmental genes that underlie morphological evolution^65^. By contrast, divergence of core clock genes might represent a more complex evolutionary trajectory, necessitating the coevolution of multiple interacting loci.

*Pdf* evidently does not explain the entirety of the species differences in plasticity: there are almost certainly contributions of additional loci that we have not tested and/or more complex genetic interactions that we cannot identify with our screen design. For example, the possibility of transvection (*trans*-regulation of alleles on homologous chromosomes)^24^ in our hybrid screen might have masked the contributions of some divergent loci. Our observation of differences between *Pdf cis*-regulatory activity using transcriptional reporters, and endogenous *Pdf* RNA and Pdf protein levels in *D. sechellia* and *D. melanogaster* suggest that additional genes might nevertheless ultimately impact Pdf. Beyond species-specific *cis*-regulation characterized here, post-transcriptional regulation of *Pdf*^31^, as well as control of the transport and secretion of this neuropeptide^66^ are also potentially subject to divergent regulation. Indeed, in *D. melanogaster*, a *D. virilis* transcriptional reporter for *Pdf* labels the s-LNvs (in addition to several non-circadian cells) despite this species, like other high-latitude drosophilids^4,16^, lacking endogenous Pdf expression in these neurons^5^; these observations suggest that mechanisms other than (or in addition to) *cis*-regulatory divergence underlie this spatial difference in expression. Finally, molecules functioning downstream of Pdf in controlling plasticity (e.g., Eyes absent^67^) are possible loci of evolutionary adaptations.

*D. sechellia* also displays greatly reduced morning activity compared to other drosophilids. As this phenotype is similar to that observed in *D. melanogaster Pdf* mutants^68^, it is perhaps surprising that we did not find an effect of the *Pdf* locus itself in this difference. Rather, we identified several genes with the ability to regulate *Pdf* expression in *trans*, highlighting a different evolutionary trajectory to divergence of circadian plasticity that nevertheless converges upon this neuropeptide. The morning and evening oscillators partially overlap in function, sharing synaptic feedback^27,69^, with both being required for long photoperiod adaptation^35^. The mechanistic and evolutionary connection (if any) between divergence of circadian plasticity and morning activity warrants further exploration. The reason, if any, for reduced morning peak activity in *D. sechellia* remains unclear; this issue might be illuminated by future analysis of the circadian pattern of other aspects of this species’ behavior, such as courtship or feeding.

*D. sechellia*’s loss of photoperiod plasticity is particularly intriguing in the context of this species’ specialist ecology for the noni fruit of *Morinda citrifolia*, on which it exclusively feeds and breeds. Noni specialization has involved substantial evolution of its chemosensory behaviors^59,70-75^. Our work extends knowledge of this species’ phenotypic divergence to non-host related behaviors. Why loss of circadian plasticity of *D. sechellia* likely leads to a severe fitness cost at high latitudes is unknown, but we speculate that longer photoperiods result in altered pheromone production, as observed in different seasonal morphs of *Drosophila suzukii^76^*. A more general consideration is why *D. sechellia* has lost circadian plasticity. One hypothesis is that in a constant environment, selection to maintain plasticity mechanisms is relaxed, leading them to degenerate over time. Alternatively, in stable environments, plasticity might come at a fitness cost, leading selection to favor its loss under constant conditions to enhance, for example, the robustness of this species’ circadian activity. While we cannot currently discriminate between these two possibilities, if the latter is correct, our view of *D. sechellia*’s specialization must expand beyond host fruit preference evolution to restriction to an equatorial environment. Indeed, *D. sechellia*’s circadian phenotype might contribute to its restriction to the Seychelles archipelago, despite the much larger modern range of *M. citrifolia*^77^. Exploration of the impact of differences in circadian plasticity mechanisms to latitudinal constraint of other species seems warranted.

## Acknowledgements

We are grateful to Enrico Bertolini, Charlotte Förster, Daniel Matute, Julia Saltz, the Bloomington *Drosophila* Stock Center (NIH P40OD018537) and the Developmental Studies Hybridoma Bank (NICHD of the NIH, University of Iowa) for reagents. We thank Roman Arguello, Enrico Bertolini, David Gatfield and members of the Benton laboratory for discussions and comments on the manuscript. Research in E.N.’s laboratory is supported by the SNSF (310030_189169), and R.B.’s laboratory is supported by the University of Lausanne, an ERC Advanced Grant (833548) and the SNSF (310030B_185377).

## Author Contributions

M.P.S. and R.B. conceived the project. M.P.S. designed, performed and analyzed most experiments. L.A. performed the hybrid screening and sequencing of the *Pdf* gene region. L.L.D.D. contributed to the experiments in Fig. 1c. J.C. contributed to the experiments in Extended Data Fig. 3 and 4. R.K. assisted with preliminary behavioral experiments and, together with E.N. and R.B., provided input on experimental design, analysis and interpretation. M.P.S. and R.B. wrote the paper with feedback from all other authors.

## Methods

### *Drosophila* strains and rearing

All flies were reared on a wheat flour-yeast-fruit juice media in non-overlapping 2-week cycles kept in 12:12 h LD at 25°C. For *D. sechellia* strains, we added an additional mixture of instant *Drosophila* medium (Formula 4-24 blue, Carolina bio-supply) mixed with juice of their host noni fruit (Raab Vitalfood).

For behavioral comparisons of *D. melanogaster*, *D. sechellia*, *D. simulans,* and *D. mauritiana* circadian behavior, at least two wild-type strains of each species were used (*Dmel*CS, *Dmel*OR, *Dmel*LZV L72 *, Dmel*LZV L76*, Dsec*07, *Dsec*28, *Dsim*MD221, *Dsim*MD242, *Dmau*90, and *Dmau*91). To screen candidate genes for effects on differences in circadian behavior between *D. melanogaster* and *D. sechellia*, we used *D. melanogaster* strains containing known loss-of-function mutations for genes previously associated with circadian behavior in *D. melanogaster*. A list of all fly strains and their hybridization success (when applicable), is provided in Supplementary Table 1. When strains were not available (*vrille*) or not hybridizable (*Jet* and *timeless*), we used *D. melanogaster* deficiency strains, containing engineered chromosomal deletions spanning the region of a candidate gene, in addition to many other loci^79^. In the case of *timeless*, a deficiency strain did not hybridize either. The *Pdf* strain we used is the *pdf^01^* allele^68^ in the Canton-S genetic background (provided by Charlotte Förster), as we were unable to hybridize the original *pdf^01^* strain. To confirm that the effect we found (Extended Data Fig. 3) was not due to a difference in *D. melanogaster* genetic background, we additionally compared it to the same parental Canton-S (denoted here *Dmel*CS^W^, where “W” = Würzburg), which displayed qualitatively similar pre-dawn activity and circadian plasticity to our own *Dmel*CS strain (Extended Data Fig. 9).

### Hybrid crosses and circadian candidate gene screening

To screen available circadian candidate genes, we created *D. melanogaster/D. sechellia* hybrids as previously described^80^. In brief, very young virgin females were crossed to males that were collected as virgins and aged in high density for 5-7 days. To increase their interactions, we pushed a plug into the vial to leave 2-3 cm height above the food surface. These crosses yield only sterile but viable males. This method does not allow us to test sex-linked candidate genes, such as the core transcriptional feedback loop member *period*^81^, and the Pdf receptor gene, *Pdfr*^64^. We aimed to phenotype at least 15 hybrids of each genotype, but due to the strong reproductive isolation between species, some genotypes were difficult to cross to *D. sechellia*, resulting in a low sample size.

### *Drosophila* activity monitoring

For all activity monitoring, we used 1-3 day old males in the *Drosophila* activity monitor (DAM) system^82^ stored in small incubators that continuously monitor light and temperature conditions (TriTech Research DT2-CIRC-TK). In brief, this system uses an infrared beam that bisects a 5 mm glass tube, in which the fly is stored, to record activity as the number of beam crosses per minute. Flies are stored in tubes with a 5% sucrose 2% agar w/v solution at one end, and capped with a cotton plug at the other. Each monitor records the activity of up to 32 flies simultaneously, and multiple monitors are stored in a single incubator. For each genotype, we recorded flies over at least two technical replicates.

All flies were first exposed to 7 days of 12:12 h LD, and then shifted to one of four extended photoperiod cycles for an additional 7 days: 14:10, 16:8, 18:6, or 20:4 h LD to allow us to measure 12:12 h LD-associated (i.e., morning anticipation) and extended photoperiod-associated behaviors (i.e., evening peak plasticity) for each fly. For assessment of free-running period, flies were exposed to 7 days of DD following 7 days of 12:12 h LD. For each photoperiod regime, we took the average activity of the final 4 days of each week-long period. The initial 3 days were considered an acclimation period, and were discarded. All subsequent analyses were performed in R using the Rethomics package^83^.

To quantify pre-dawn activity, the average normalized activity was calculated for each fly in 30 min bins in the 3 h preceding dawn. To quantify morning and evening peak times, peak activity was identified from the average activity of each fly in 10 min bins during the last 4 days of both the 12:12 h LD and extended photoperiod using custom R scripts (available at: github.com/mshahandeh/circ_plasticity). First, a rolling triangular mean was applied to smooth the data. The data was split into two 12 h sections, the first spanning the time around lights-on and the second spanning the time around lights-off (at least 3 h preceding and 3 h after for both). The global peak was identified within each data set and recorded as the timing of the morning peak and evening peak, respectively.

### Construction of transgenic lines

∼2.4 kb upstream of the *Pdf* start codon was PCR amplified from *D. melanogaster* (*Dmel*CS) or *D. sechellia* (*Dsec*28) genomic DNA and Gateway cloned into the pDONR221 vector, sequenced-verified, and subcloned into both pHemmarG (Addgene #31221) for CD4:tdGFP reporters, and pBPGUw (Addgene #17575) for Gal4 drivers. Constructs were injected and integrated into the attP2 landing site (chromosome 3) in the *D. melanogaster* genome by BestGene Inc. Oligonucleotides used for cloning and sequence verification are listed in Supplementary Table 3.

### Single molecule mRNA FISH

We performed single molecule mRNA fluorescent in-situ hybridization to quantify *pdf* mRNA expression at various time points in the s-LNvs and l-LNvs. We used the *Pdf* probe library described previously^45^ bound to the Cy5 fluorophore (LubioScience), and adapted a published protocol^84^. Brains were imaged using an inverted confocal microscope (Zeiss LSM 710 or 880) equipped with a 40× or 63× oil immersion objective using fixed settings to maximize comparability of images within experiments. Images were processed in Fiji and RNA spots were counted using the Fiji macro RS-FISH^85^. No signal was detected outside of the LNv cell bodies. We compared RNA spot counts between strains within photoperiod treatments using a Wilcoxon rank-sum test followed by post-hoc correction for multiple tests^86^. We did not compare between experiments as these flies were dissected, stained and imaged separately. We repeated smFISH throughout the morning peak to ensure replicability of the overall pattern of expression. We did not pool these data as they are from a separate staining/imaging and may not be as comparable.

### Immunofluorescence

For immunofluorescence of whole-mount *Drosophila* brains, 1-2 day old males were collected and acclimated to a specific photoperiod for 4 additional days. To standardize sampling times, we fixed these flies in 4% paraformaldehyde for 2 h at room temperature with gentle agitation prior to dissection. Brains were dissected and stained essentially as described^87^. Primary and secondary antibodies and concentrations used are provided in Supplementary Table 2. Brains were imaged using an inverted confocal microscope (Zeiss LSM 710 or 880) equipped with a 20× or 40× objective using fixed settings to maximize comparability of images. To quantify fluorescence, images were processed in Fiji, by first creating a maximum intensity projection Z-stack which was then thresholded to remove background signal^88^. Relative fluorescence was measured for each set of neurons by structure (i.e., LNv soma or s-LNv dorsal axonal projections) as integrated density of pixel intensity, and the average of both hemispheres was recorded for each brain. We quantified all images blind to treatment (species, genotype, and timepoint). We compared Pdf immunofluorescence between strains within photoperiod treatments using a Wilcoxon rank-sum test followed by post-hoc correction for multiple tests^86^. We did not compare between experiments as these flies were dissected, stained, and imaged separately. We repeated immunostainings throughout the morning peak to ensure replicability of the overall pattern of expression. These data cannot be pooled, however, as they are from a separate staining/imaging and produce different fluorescence measurements (arbitrary units).

To compare structural plasticity of s-LNv axonal projections between *D. melanogaster* and *D. sechellia*, we imaged the most dorsal projections during two timepoints in the light and dark phase (2 h and 14 h, respectively) at 40× with a 2× digital zoom. We performed a Scholl analysis, counting the number of axonal crossings with concentric 10 µm arcs using the Neuroanatomy Fiji plugin^89^. The number of axonal crossings was averaged per hemisphere for each brain and compared using a Wilcoxon rank-sum test. We performed this experiment in two replicates and pooled replicates for analysis as fluorescence intensity was not measured.

### *Pdf* gene region sequence comparisons

We Sanger-sequenced the *Pdf* gene region using the oligonucleotides listed in Supplementary Table 3. The sequences were assembled and aligned in SnapGene software (www.snapgene.com) using MUSCLE v3.8.1551^90^ and then visually inspected for errors. When appropriate, sequences were translated to an amino acid alignment and visualized using Jalview^91^. For 5’-regulatory sequences, we used the R package phangorn to generate maximum likelihood trees^92^, using the modelTest function to identify the best fitting substitution model and performing standard bootstrapping to obtain support values. The MEME program was used to discover putative regulatory motifs common across all sequences^47^. We restricted this analysis to the top ten significant motifs identified.

### Population genetic analysis of the *Pdf* 5’ regulatory sequences in *D. melanogaster* and *D. sechellia*

To detect genomic patterns of clinal adaptation in the *Pdf* 5’ sequences, we used a dataset of single nucleotide variants in globally distributed *D. melanogaster* populations^48^. We calculated the average MAF for each population in this region, and for the same-sized region upstream of the start codon of 6 control neuropeptide genes. Spearman’s rho was used to correlate MAF with the latitude of the capital city in each country where the populations were sampled (precise latitudes were not available).

### Longevity assay

To test for photoperiod-dependent differences in lifespan, we acclimated 1-day old *Dmel*CS, *Dmel*OR, *Dsec*07 and *Dsec*28 males to either 12:12 h LD or 16:8 h LD conditions. We held these flies in vials containing wheat flour-yeast-fruit juice media to which we added an additional mixture of instant *Drosophila* medium (Formula 4-24 blue, Carolina bio-supply) mixed with noni juice for *D. sechellia* and apple cider vinegar for *D. melanogaster*. Flies were transferred to fresh vials every 3 days to prevent the media from drying out. We recorded for each week-day the number of vials in which a fly died until all flies among one treatment per strain were dead. No significant differences were detected between strains within species (Fisher’s exact test, all p > 0.05), which were therefore pooled to represent the species for analysis. We compared cumulative survival probability using the R package ‘survival’^93^.

### Copulation rate assays

To test for photoperiod-dependent differences in copulation rate, we first acclimated 1-day old virgin *Dmel*CS, *Dmel*OR, *Dsec*07 and *Dsec*28 males and females to either 12:12 h LD or 16:8 h LD conditions for 4 days. For our short-term assay, we aspirated single females into 25 mm food vials containing wheat flour-yeast-fruit juice media, returned them to their respective photoperiods, and allowed them to recover for 24 h. The following day, 30 min after lights-on, we aspirated a single male of the same genotype into each tube, pushed the plug into the vial so that pairs had 2 cm above the food surface, forcing them to interact. We observed for copulation for 2 h, recording successfully and unsuccessfully copulating pairs. For our long-term assay, we similarly acclimated flies for 4 days, but aspirated male-female pairs of the same genotype into vials and returned them to their respective photoperiods. We transferred these pairs to new vials every 24 h, and scored copulation success per day based on the presence of offspring. Flies that produced no offspring over 7 days were considered to have never mated. We observed no differences between strains within species in either experiment (Fisher’s exact test, all p = 1), so these data were pooled to represent the species for analysis. Copulation frequencies within species between treatments were compared using a Wilcoxon rank-sum test.

---

**Extended Data Fig. 1.**
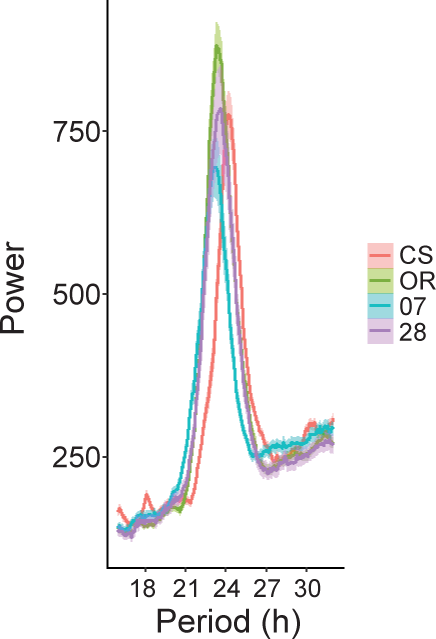
*D. melanogaster* and *D. sechellia* strains exhibit ∼24 h periods. Periodogram analysis from 5 days of constant darkness (DD) for *D. melanogaster* (CS and OR) and *D. sechellia* (07 and 28) strains. Period estimates: CS (24.36 h), OR (23.45 h), 07 (23.16 h), 28 (23.57 h). Sample sizes as in Fig. 1e

**Extended Data Fig. 2.**
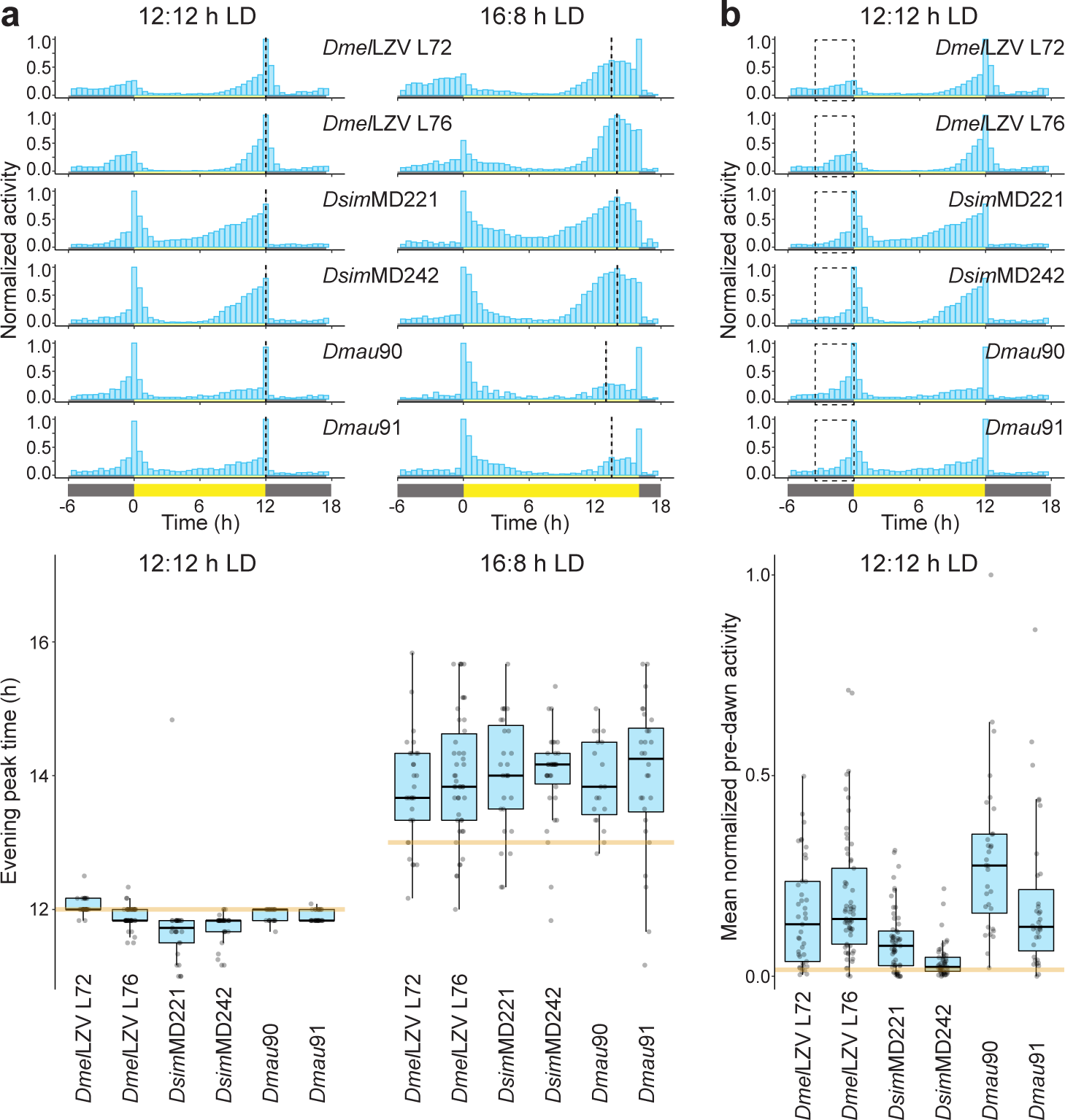
Tropical *D. melanogaster*, *D. simulans* and *D. mauritiana* strains display prominent circadian plasticity and morning anticipation. **a**, Top: mean normalized activity of two recently-collected strains of *D. melanogaster* (LZV L72 and LZV L76, from the Lower Zambezi Valley), *D. simulans* (MD221 and MD242, from Madagascar) and *D. mauritiana* (*Dmau*90 and *Dmau*91, from Mauritius) under the indicated photoperiods. Plots depict average activity of the last 4 days of a 7-day recording period. Dashed lines highlight the average evening peak time. Bottom: evening peak time for these flies. The orange line depicts the median evening peak time of individuals of both *D. sechellia* strains (from Fig. 1c). Sample sizes as follows: LZV L72 (29), LZV L74 (46), MD221 (27), MD242(34), *Dmau*90 (19), *Dmau*91 (28). **b**, Top: mean normalized activity of the same strains as in **a** under a 12:12 h LD cycle (same as in **a**). Plots depict average activity of the last 4 days of a 7-day recording period. Dashed boxes highlight the pre-dawn period, 3 h before lights-on. Bottom: Mean normalized activity of individual flies within this pre-dawn period. Sample sizes as follows: LZV L72 (41), LZV L74 (61), MD221 (57), MD242 (52), *Dmau*90 (33), *Dmau*91 (34). The orange line depicts the median pre-dawn activity of individuals of both *D. sechellia* strains (from Fig. 1d).

**Extended Data Fig. 3.**
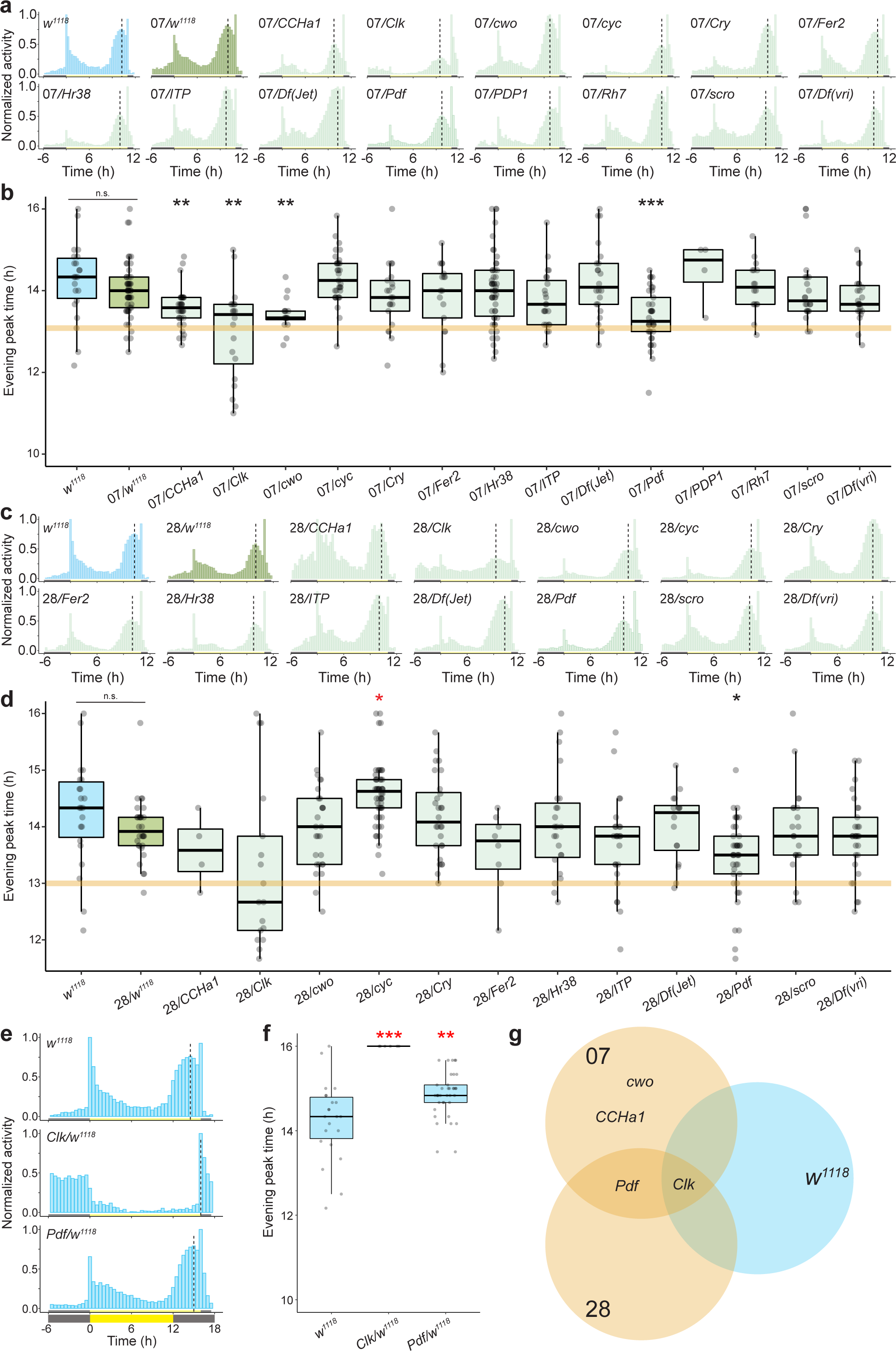
Screen results for the genetic basis of interspecific differences in circadian plasticity. **a**, Mean normalized activity of the indicated control and hybrid genotypes under a 16:8 h LD cycle. Plots depict average activity of the last 4 days of a 7-day extended photoperiod, following 7 days of 12:12 h LD. Vertical dashed lines indicate the average timing of the evening peak for each strain. Sample sizes as follows: *w^1118^* (22), 07/*w^1118^* (53), 07/*CCHa1* (34), 07/*Clk* (20), 07/*cwo* (16), 07/*cyc* (33), 07/*Cry* (21), 07/*Fer2* (17), 07/*Hr38* (50), 07/*ITP* (23), 07/*Jet* (22), 07/*Pdf* (37), 07/*PDP1* (4), 07/*Rh7* (16), 07/*scro* (22), 07/*vri* (23). **b**, Evening peak time for the flies depicted in **a**. Asterisks indicate significant differences: ** p < 0.01 and *** = p < 0.001 (Wilcoxon tests comparing each test hybrid to the control hybrid strain (07/w^1118^) with Bonferroni correction). n.s. = not significantly different. The orange line marks the median evening peak delay of the *D. sechellia* parental strain (07). **c**, Mean normalized activity of the indicated control and hybrid genotypes under a 16:8 h LD cycle. Plots depict average activity of the last 4 days of a 7-day extended photoperiod, following 7 days of 12:12 h LD. Vertical dashed lines indicate the average timing of the evening peak for each strain. Sample sizes as follows: *w^1118^* (22), 28/*w^1118^* (31), 28/*CCHa1* (4), 28/*Clk* (17), 28/*cwo* (27), 28/*cyc* (52), 28/*Cry* (28), 28/*Fer* (8), 28/*Hr38* (23), 28/*Itp* (25), 28/*Jet* (16), 28/*Pdf* (40), 28/*scro* (31), 28/*vri* (29), 28 (19). **d**, Evening peak time for the flies depicted in **c**. Asterisks indicate significant differences: * = p < 0.05 (Wilcoxon tests comparing each test hybrid to the control hybrid strain (28/*w^1118^*) with Bonferroni correction). Red asterisks denote a significant increase in circadian plasticity. n.s. = not significantly different. The orange line marks the median evening peak delay of the *D. sechellia* parental strain (28). **e**, Mean normalized activity of the indicated hemizygous *D. melanogaster* genotypes that displayed an effect in both hybrid backgrounds under a 16:8 h LD cycle. Plots depict average activity of the last 4 days of a 7-day extended photoperiod, following 7 days of 12:12 h LD. Vertical dashed lines indicate the average timing of the evening peak for each strain. Sample sizes as follows: *w^1118^* (22), *Clk*/*w^1118^* (6), *Pdf*/*w^1118^* (37). **f**, Evening peak time for the flies depicted in **e**. Asterisks indicate significant differences: *** = p < 0.001 and ** = p < 0.01 (Wilcoxon tests comparing each test hemizygote to the control strain (*w^1118^*) with Bonferroni correction). Red asterisks denote a significant increase in circadian plasticity. **g**, Summary of the overlapping hits. *A priori*, we considered the strongest candidates would display a reduction in circadian plasticity in both *Dsec*07 and *Dsec*28 hybrids, but not in *w^1118^* hemizygotes; only *Pdf* fulfilled these criteria. Note: the *Clk* mutant used in this screen is a dominant negative allele, and thus we expect the behaviour of *Clk/w^1118^* mutants to display a total *Clk* loss-of-function phenotype. Interestingly, we do not observe this phenotype in either test hybrid genotype, indicating divergence of the *Clk* locus between species. If and how this divergence affects behaviour requires subsequent investigation.

**Extended Data Fig. 4.**
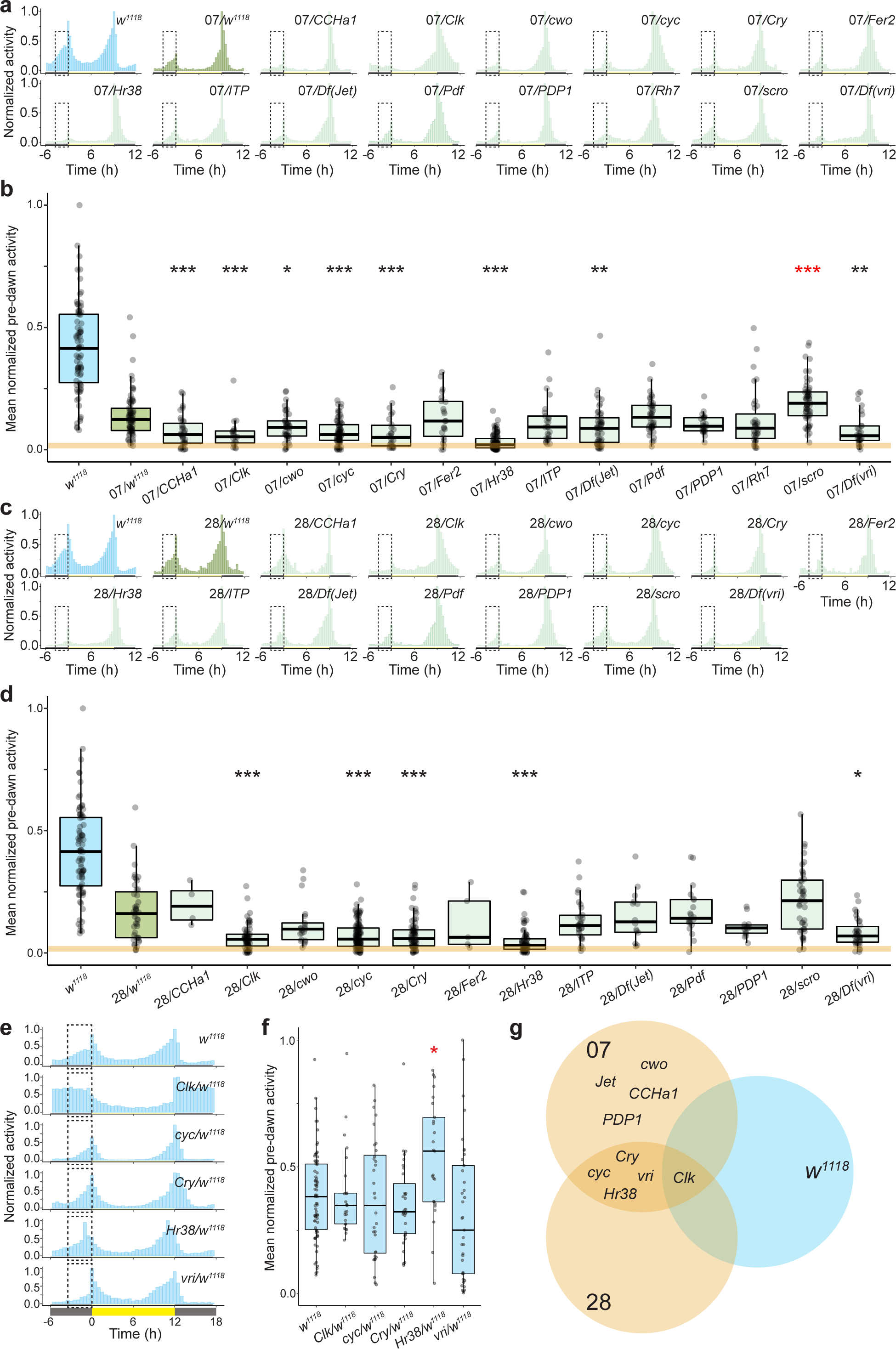
Screen results for the genetic basis of interspecific differences in morning anticipation. **a**, Mean normalized activity of the indicated genotypes under a 12:12 h LD cycle. Dashed boxes highlight the pre-dawn area used to quantify morning anticipation. Sample sizes as follows: *w^1118^* (78), 07/*w^1118^* (69), 07/*CCHa1* (21), 07/*Clk* (19), 07/*cwo* (43), 07/*cyc* (43), 07/*Cry* (26), 07/*Fer* (18), 07/*Hr38* (87), 07/*ITP* (23), 07/*Jet* (49), 07/*Pdf* (42), 07/*PDP1* (23), 07/*Rh7* (33),07/*scro* (58), 07/*vri* (28), 07 (40). **b**, Mean normalized pre-dawn activity for the genotypes in A. Asterisks indicate significant differences: ** = p < 0.01 and *** = p < 0.001 (Wilcoxon tests comparing each test hybrid to the control hybrid strain (07/*w^1118^*) with Bonferroni correction). Red asterisks denote a significant increase in circadian plasticity. The orange line marks the median pre-dawn activity of the *D. sechellia* parental strain (07). **c**, Mean normalized activity of the indicated genotypes under a 12:12 h LD cycle. Dashed boxes highlight the pre-dawn area used to quantify morning anticipation. Sample sizes as follows: w^1118^ (78), 28/w^1118^ (22), 28/*CCHa1* (4), 28/*Clk* (64), 28/*cwo* (20), 28/*cyc* (56), 28/*Cry* (66), 28/*Fer* (5), 28/*Hr38* (43), 28/*ITP* (25), 28/*Jet* (22), 28/*Pdf* (21), 28/*PDP1* (14), 28/*scro* (38), 28/*vri* (33), 28 (36). **d**, Mean normalized pre-dawn activity for the genotypes in C. Asterisks indicate significant differences: * = p < 0.05 and *** = p < 0.001 (Wilcoxon tests comparing each test hybrid to the control hybrid strain (28/w^1118^) with Bonferroni correction). The orange line marks the median pre-dawn activity of the *D. sechellia* parental strain (28). **e**, Mean normalized activity of the indicated hemizygous *D. melanogaster* genotypes that displayed an effect in a hybrid background under a 12:12 h LD cycle. Dashed boxes highlight the pre-dawn area used to quantify morning anticipation. (07 or 28) under a 12h LD cycle. Plots depict average activity of the last 4 days of a 7 day recording period. Dashed boxes highlight the pre-dawn period, 3 h before lights-on. Sample sizes as follows: w^1118^ (78), *Clk*/w^1118^ (25), *cyc*/w^1118^ (31), *Cry*/w^1118^ (32), *Hr38*/w^1118^ (25), *vri*/w^1118^ (46). **f**, Mean normalized pre-dawn activity for the genotypes in **e**. Asterisks indicate significant differences: * = p < 0.05 (Wilcoxon tests comparing each test hemizygote to the control strain (*w^1118^*) with Bonferroni correction). Red asterisks denote a significant increase in pre-dawn activity. **g**, Summary of the overlapping hits from each of the above genotypes. *A priori*, we considered the strongest candidates to display a reduction in morning anticipation in *Dsec*07 and *Dsec*28 hybrids, but not in *w^1118^* hemizygotes. See Extended Data Fig. 3g legend for notes on the *Clk/w^1118^* mutant phenotype.

**Extended Data Fig. 5.**
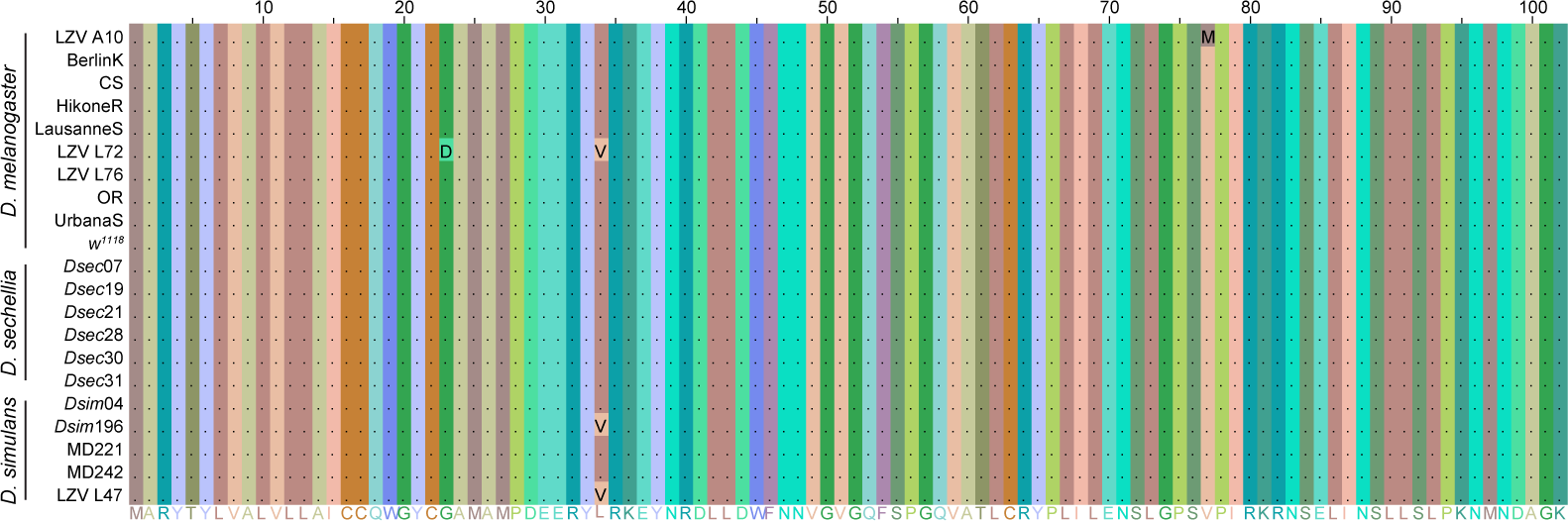
The predicted Pdf protein sequence is highly conserved between *D. melanogaster*, *D. sechellia* and *D. simulans*. Alignment of the predicted Pdf protein sequence of 10 *D. melanogaster*, 6 *D. sechellia* and 5 *D. simulans* strains. Amino acid residues are coloured by similarity, periods indicate conserved amino acid residues and letters indicate variable residues. No fixed differences are observed between species. The consensus sequence is displayed at the bottom.

**Extended Data Fig. 6.**
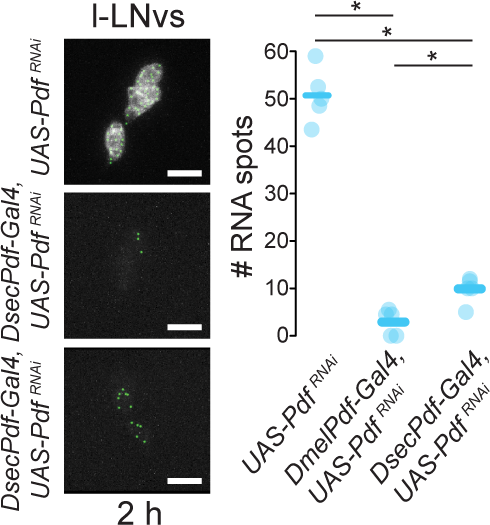
Validation of differential *Pdf* transcript depletion by smFISH. Left: representative smFISH images for one genetic control (*UAS-Pdf^RNAi^/+*), *DmelPdf-Gal4/UAS-Pdf^RNAi^* and *DsecPdf-Gal4/UAS-Pdf^RNAi^* strains with RNA spots identified by RS-FISH. Right: RNA spot quantifications. N = 5 for each genotype.

**Extended Data Fig. 7.**
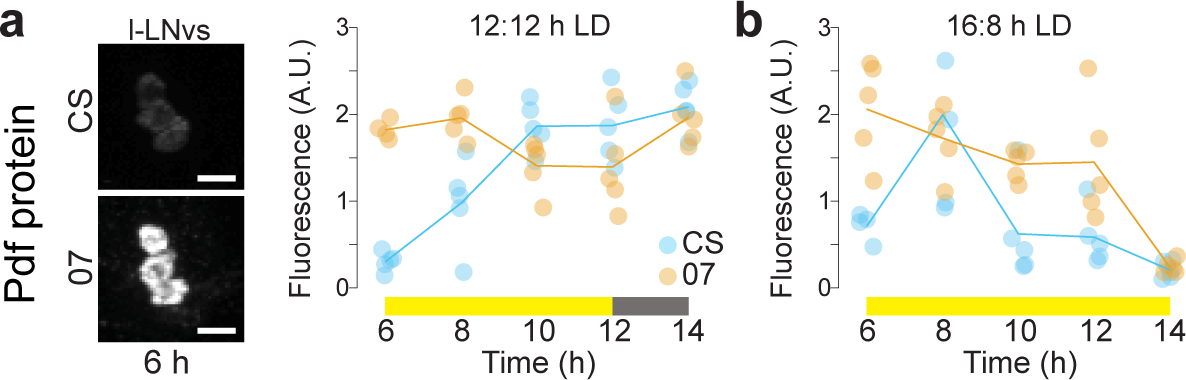
Pdf immunofluorescence in the l-LNvs of *D. melanogaster* and *D. sechellia* during the evening peak. **a**, Left: representative images of Pdf immunofluorescence in the l-LNv soma for the CS and 07 strains under 12:12 h LD at one time point (6 h). Right: quantifications of Pdf signals at 5 time points spanning the evening activity peak period. **b**, Quantifications of Pdf signals at 5 time points spanning the evening activity peak period in the l-LNv soma for the CS and 07 strains under 16:8 h LD.

**Extended Data Fig. 8.**
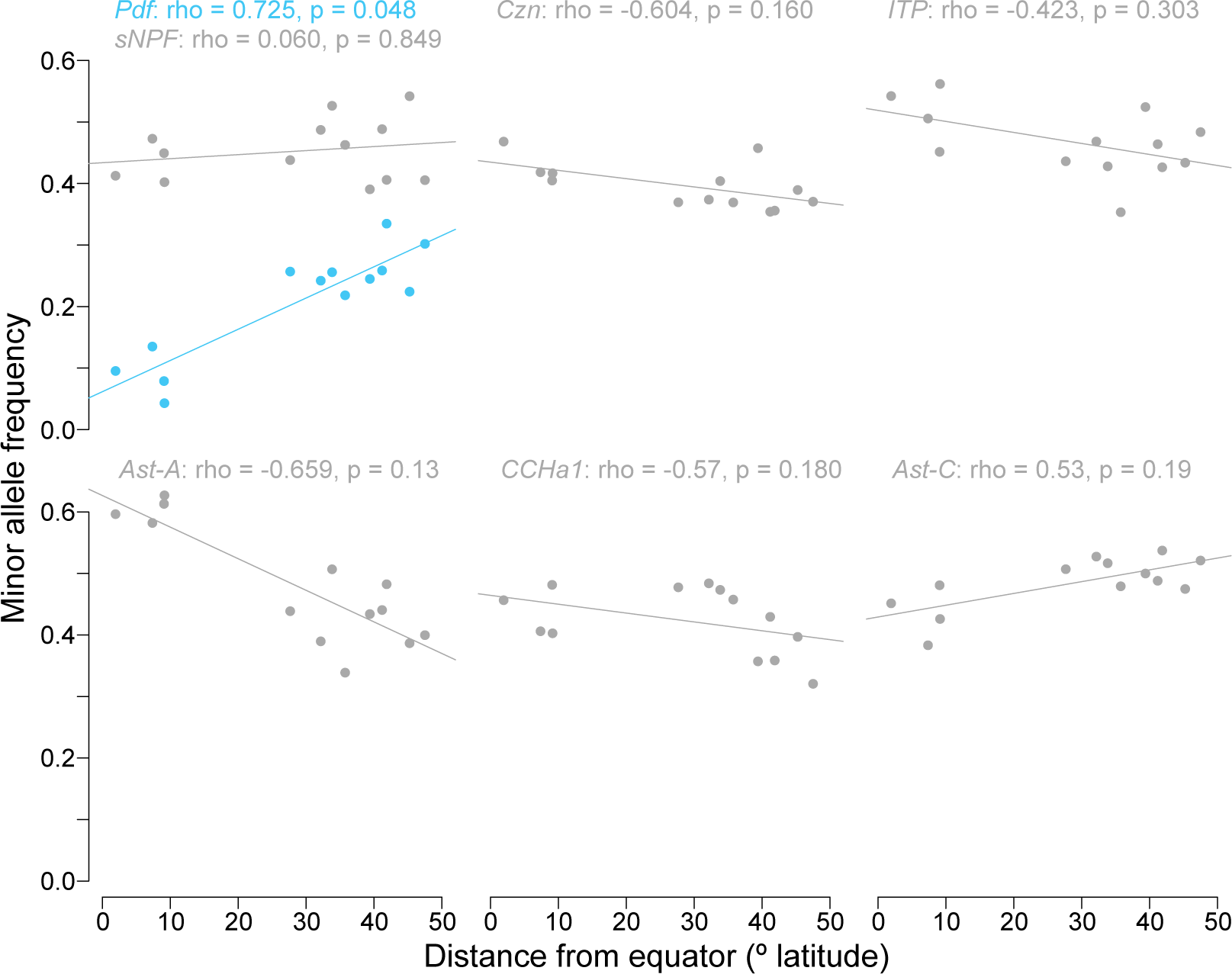
Correlations of minor allele frequency and latitude in *D. melanogaster* populations for control neuropeptide genes. Each plot depicts the correlation between minor allele frequency and latitude for the putative 5’-regulatory region (∼2.4 kb upstream of the start codon) for the indicated neuropeptide genes. For reference, the analysis for the *Pdf* 5’-regulatory region (from Fig. 5c) is shown on the first plot. Values for Spearman’s rho and Bonferroni corrected p-values are listed above each plot.

**Extended Data Fig. 9.**
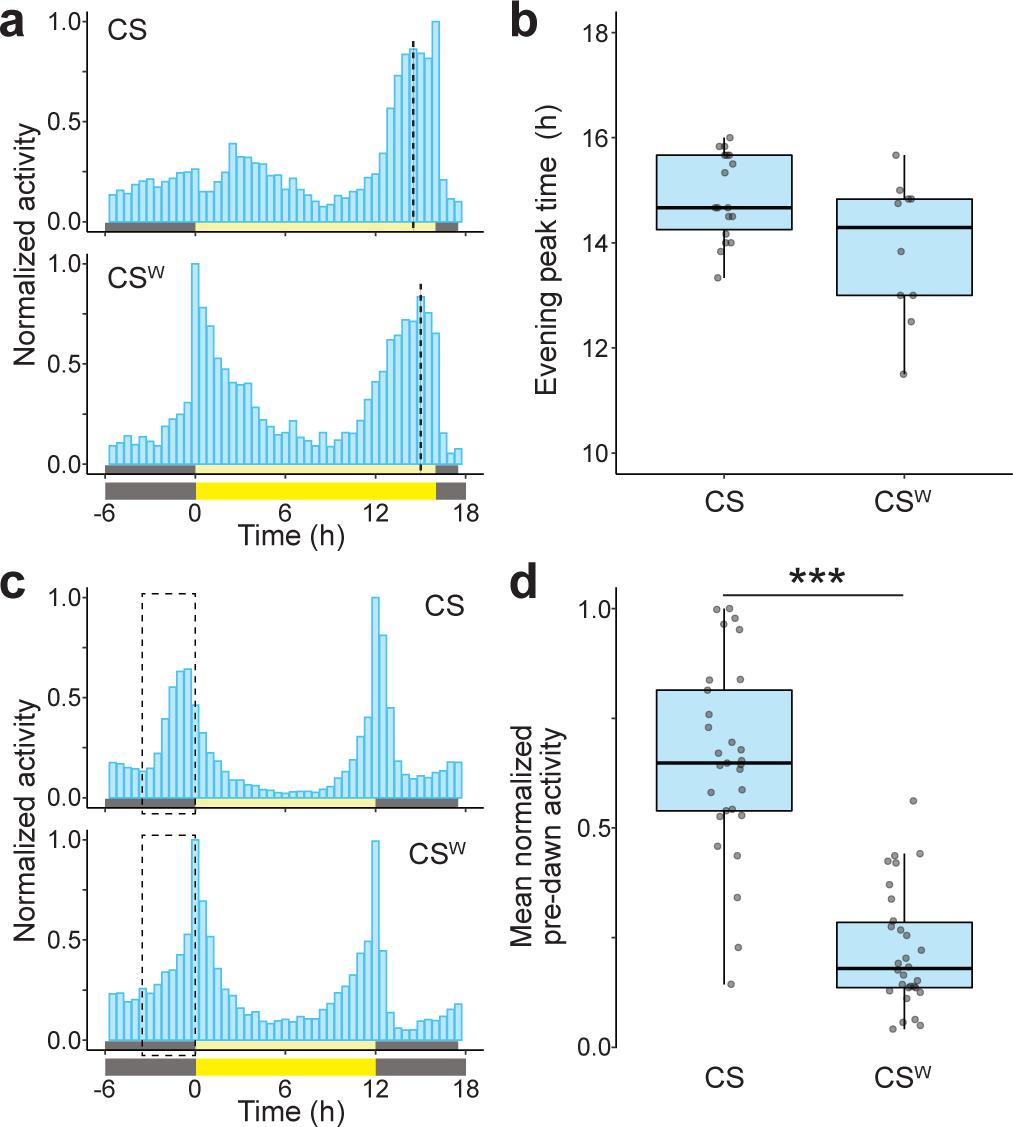
Qualitatively similar circadian plasticity and pre-dawn activity in two Canton-S strains. **a**, Mean normalized activity of two *Canton-S* (CS and CS^W^) strains collected under a 16:8 h LD cycle. Plots depict average activity of the last 4 days of a 7-day extended photoperiod, following 7 days of 12:12 h LD. Vertical dashed lines indicate the average timing of the evening peak for each strain. Sample sizes as follows: CS (18),CS^W^ (16). **b**, Evening peak time for the flies depicted in **a** is shown for each strain. No significant difference was observed between strains (Wilcoxon test). **c**, Mean normalized activity of two Canton-S strains (CS and CS^W^) under a 12:12 h LD cycle. Plots depict average activity of the last 4 days of a 7-day recording period. Dashed boxes highlight the pre-dawn period, 3 h before lights-on. Sample sizes as follows: CS (29), CS^W^ (42). **d**, The average activity for flies shown in **c** within the previously indicated pre-dawn period, mean normalized pre-dawn activity, is shown for each strain. Asterisks indicate significant differences: ** = p < 0.01 (Wilcoxon test).

## Supplementary Tables

**Supplementary Table 1.**
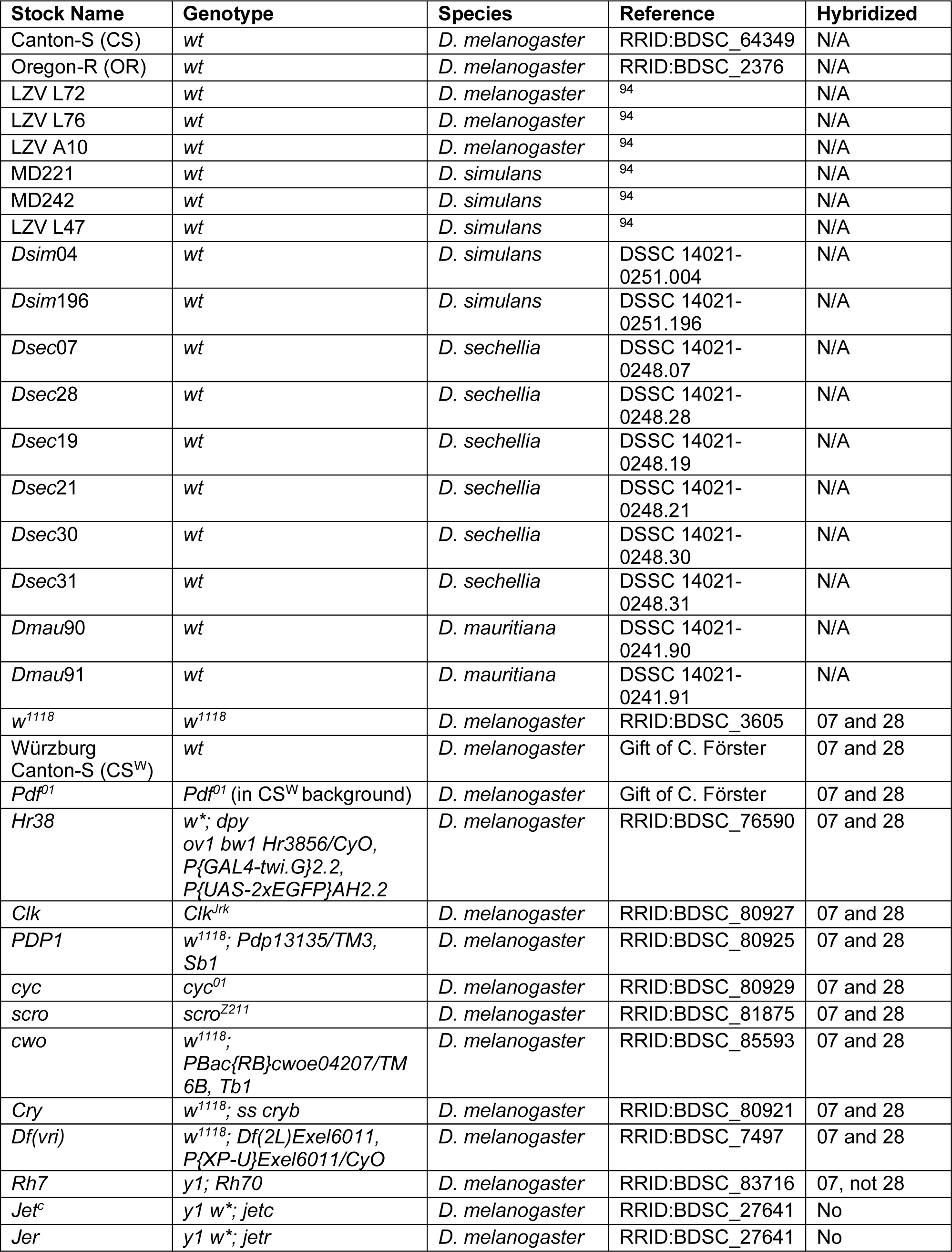
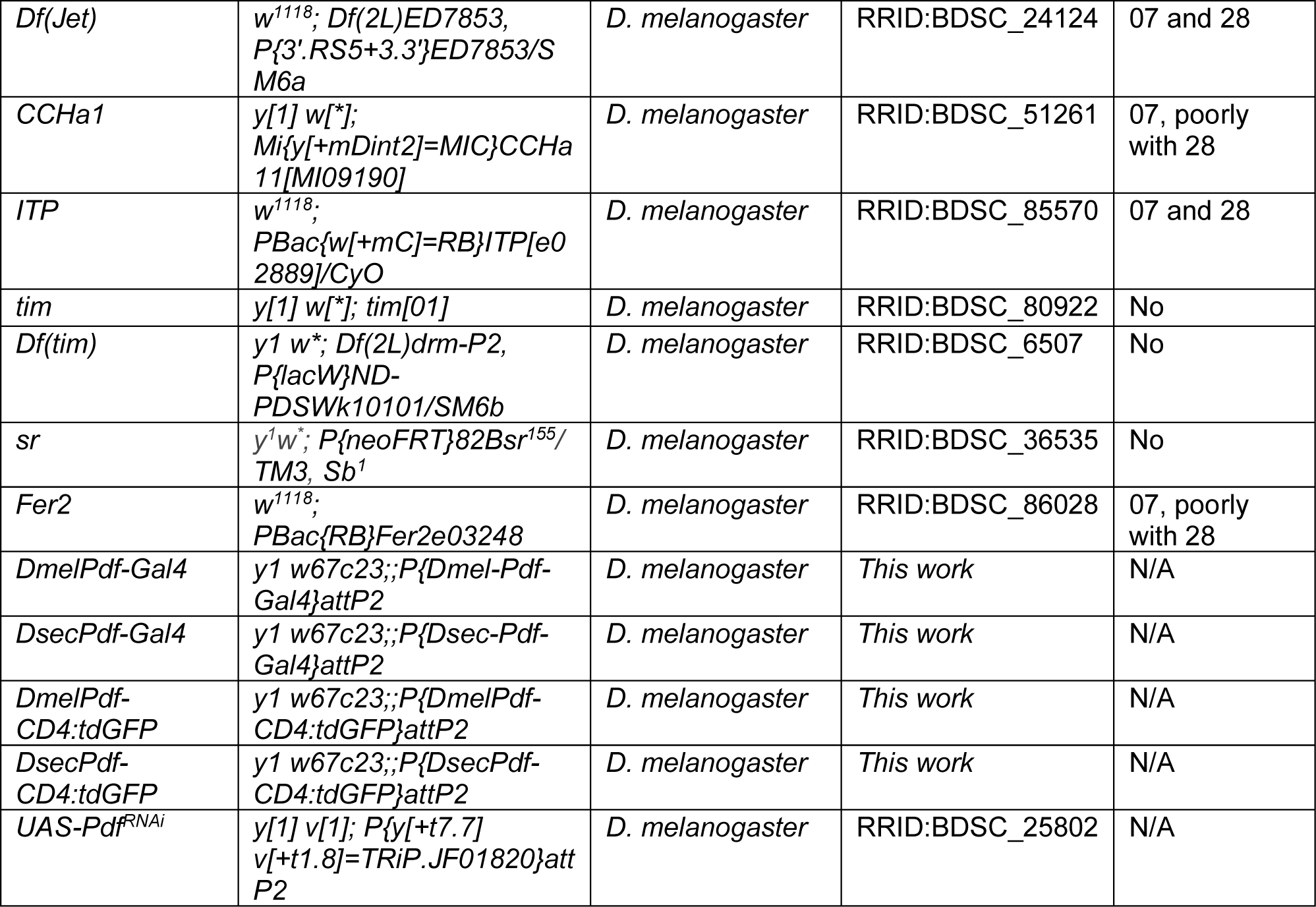
*Drosophila* strains.

**Supplementary Table 2.**
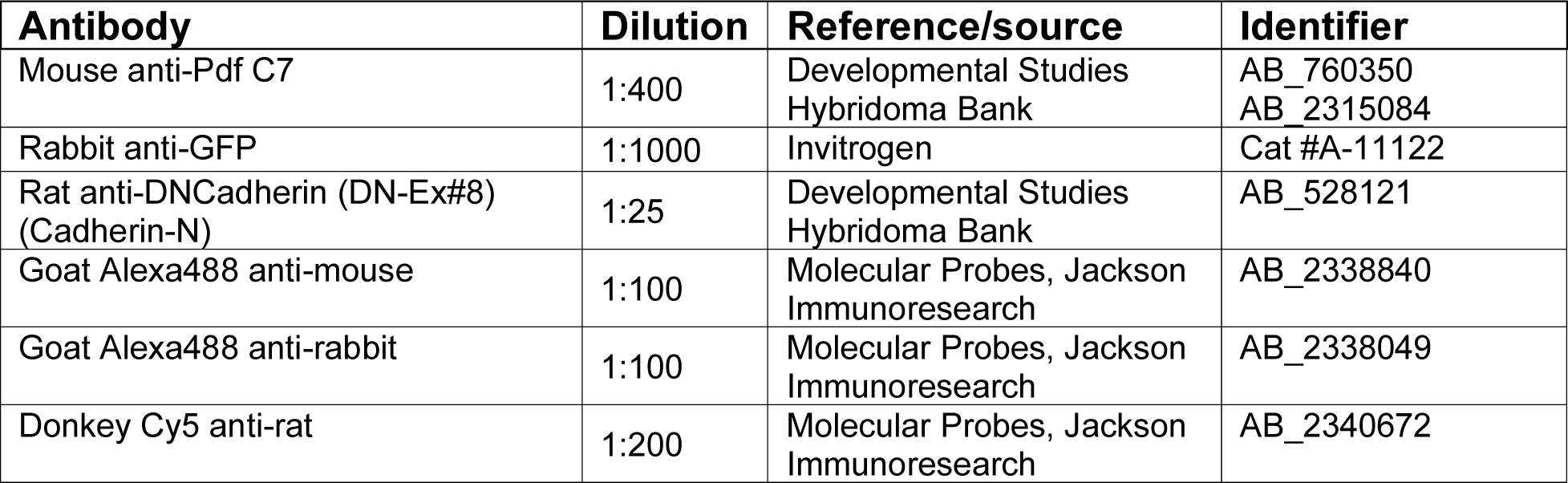
Antibodies.

**Supplementary Table 3.**
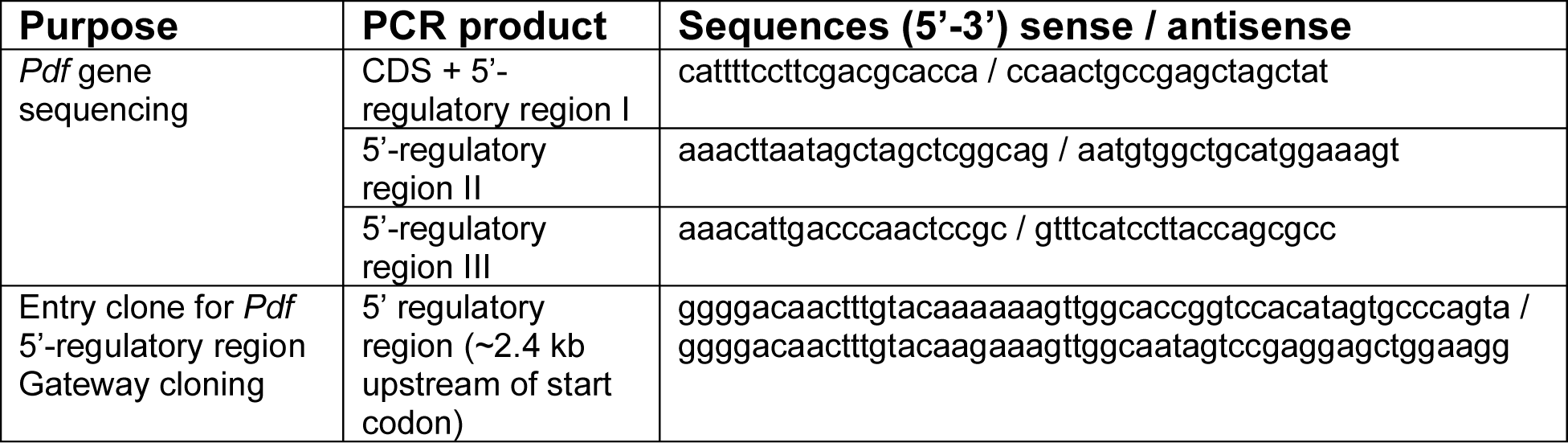
Oligonucleotides.

